# Graded sensitivity to structure and meaning throughout the human language network

**DOI:** 10.1101/2021.11.12.467812

**Authors:** Cory Shain, Hope Kean, Colton Casto, Benjamin Lipkin, Josef Affourtit, Matthew Siegelman, Francis Mollica, Evelina Fedorenko

## Abstract

Human language has a remarkable capacity to encode complex ideas. This capacity arises because language is *compositional*: the form and arrangement of words in sentences (structure) determine the conceptual relations that hold between the words’ referents (meaning). A foundational question in human cognition is whether the brain regions that support language are similarly factored into structure-selective and meaning-selective areas. In an influential study, Pallier et al. (2011, PNAS) used fMRI to investigate the brain response to sequences of real words and pseudowords and reported a sharp dissociation between structure-selective and meaning-selective brain regions. In the present study, we argue that no such dissociation emerges when individual differences in brain anatomy are considered. We report three experiments (including a close conceptual replication of Pallier et al.’s original study) that use precision fMRI methods to capture separation or overlap of function in the brains of individual participants. Our results replicate Pallier et al.’s finding that the brain’s response is modulated by the sequential structure of language but paint a different picture with respect to the structure-meaning relationship. Instead of distinct structure-selective and meaning-selective brain areas, we find distributed sensitivity to both linguistic structure and meaning throughout a broad frontotemporal brain network. Our results join a growing body of evidence for an integrated network for language in the human brain within which internal specialization is primarily a matter of degree rather than kind, in contrast with influential proposals that advocate distinct specialization of different brain areas for different types of linguistic functions.

**Significance Statement:** Using fMRI, we show that a broad network of frontal and temporal areas in the left hemisphere of the human brain is sensitive to both the structure of language and the meaning that it encodes. This finding challenges many current theories of the neurobiology of language, which propose a sharp separation between areas that encode structure and areas that encode meaning. Instead, results support a broad distribution of word- and sentence-level processing across an integrated brain network for language.

**This PDF file includes:**

Main Text

Figures 1 to 3

Tables 1 to 1

## Introduction

Human language is a powerful medium for communicating complex thoughts. This power comes from its compositional structure (1): meaning is encoded not only by individual words, but by the form and sequential arrangement of those words, which express the relationships that hold between the words’ referents. For example, the sentence *There are octopuses inside the bathtub!* is (probably) unfamiliar to the reader and also (probably) expresses a meaning with which the reader has no direct experience. Yet novel meanings are recoverable from novel sentences thanks to the systematic relationship between a sentence’s structure and its meaning. This principle even extends to unfamiliar words: when we read *There are blickets inside the dax!*, we can infer that the blickets and the dax are in a containment relationship and have certain other properties (e.g., a *blicket* is countable and a *dax* can contain something), even if we do not know the meanings of the words themselves. Thus, the expressive power of language derives from its factorization of meaning (semantics) into word-level (lexical) and compositional (combinatorial) dimensions, which are mediated by the sentence’s structure (syntax).

Many models of the neurobiology of language posit a similar factorization at the level of brain areas, such that some areas are “syntactic hubs” that selectively represent the structure of sentences, whereas others are “semantic hubs” that selectively represent the meaning of words and/or sentences, albeit with disagreement as to the precise locations of these functions in the brain (2–7). If true, this view would have fundamental implications for the organization and evolutionary origins of human cognition: brain circuits for abstract combinatorics could be recruited in service of other cognitive functions (e.g., mathematics, music, and action planning) with similar hierarchical structure to language (8–10), and they may find their origins in changes to brain anatomy that enabled algebraic thought, which were later co-opted in service of language (11, 12). One important source of evidence in favor of this view has been a landmark study by Pallier, Devauchelle, and Dehaene (ref. (13), henceforth *PDD*), who argued based on fMRI evidence for a dissociation between brain areas that selectively represent syntax and areas that selectively represent lexical (word-level) and combinatorial (sentence-level) semantics. As of this writing, PDD has been cited over 600 times, and its claims have informed theories of cognition, brain function, and evolution, especially those that posit neural circuits dedicated to abstract combinatorics (e.g., refs. (11, 12, 14–17)).

In PDD’s paradigm (**Figure 1A**), participants read 12-word stimuli presented one word at a time. These stimuli were internally composed of “chunks” (our terminology) of locally coherent connected words that varied parametrically in length. At one extreme, a stimulus contained twelve concatenated (1-word) chunks (condition “c01” in **Figure 1A**), and at the other, a stimulus contained a single 12-word chunk (condition “c12” in **Figure 1A**). In the intermediate conditions, the stimuli contained concatenated chunks of different lengths: six 2-word chunks (c02), four 3-word chunks (c03), three 4-word chunks (c04), or two 6-word chunks (c06). The chunks in these conditions always formed valid syntactic *constituents*, that is, a sequence of words dominated by a node in a tree representation of the sentence’s grammatical structure (see **Figure 1B**). PDD hypothesized that language processing requires the comprehender to maintain an increasingly complex representation of constituent structure as each new word is processed, and that this increased representational complexity will correspond to an increase in overall neuronal activity in conditions with longer constituents (because longer constituents contain more internal structure, in the form of additional constituents hierarchically nested within them; see **Figure 1B**). To investigate the abstractness of syntactic representations, a ‘Jabberwocky’ version of each condition (e.g., jab-c01, jab-c12) was created by replacing the content words (nouns, verbs, adjectives, and adverbs) with word-like nonwords (pseudowords), but preserving the syntactic ‘frame’, i.e., function words like articles and auxiliaries, and functional morphological endings (e.g., *higher and higher prices* > *hisker and hisker cleeces*).

**Figure 1.**
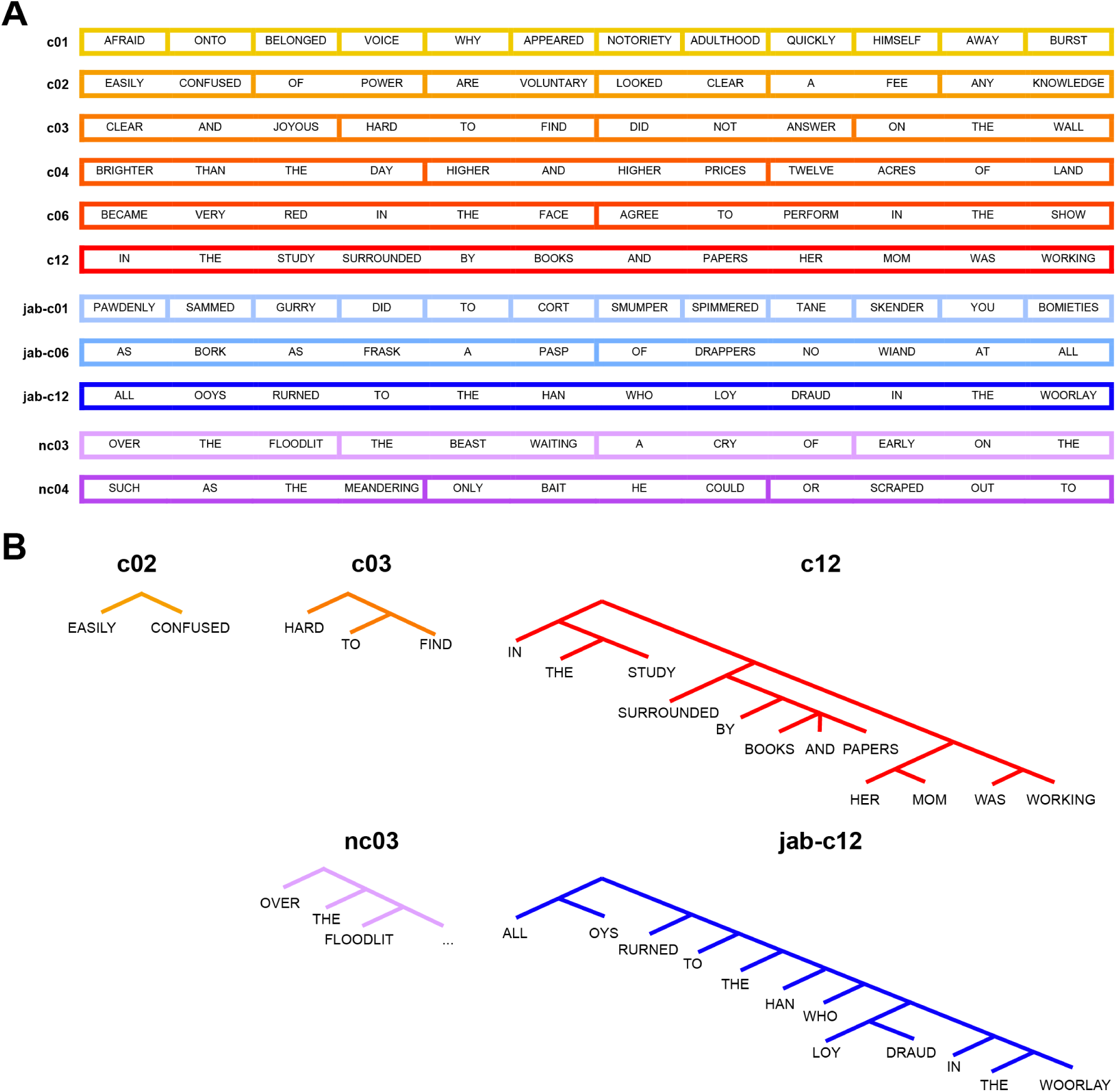
A. Examples of stimuli across conditions (from 1-word chunks, c01, to 12-word chunks, c12), with real-word constituent conditions shown in warm colors, Jabberwocky constituent conditions shown in blues, and real-word non-constituent conditions shown in purples. **B.** Visualization of constituent structure of representative chunks. In a phrase-structure grammar, a *constituent* is the entire sequence of words that is dominated by a branching node in the tree. In the c02 condition, there is exactly one constituent (“easily confused”), whereas in the c12 condition, many constituents are nested (e.g., “in the study” is a constituent nested within the entire sentence, which is itself a constituent). The same kind of nested constituency structure is implicit in the Jabberwocky condition (jab-c12), even though most of the words are meaningless. By contrast, in the non-constituent condition (nc03), the three words (“over the floodlit”) do not form a constituent, because the only node in the tree that dominates all of them (the top-most node) implicitly contains at least one additional missing word (the noun modified by “floodlit”).

This design targets three potentially dissociable dimensions of linguistic representation, each of which could be either present or absent in a given brain region’s response: **lexical semantics** (stored word meanings, which are only present in the real-word conditions), **syntax** (the implicit structure of the sentence as reflected in the forms and sequential ordering of words, which is present in both the real-word and the Jabberwocky conditions, and which increases in complexity with chunk length), and **combinatorial semantics** (the composite meaning denoted by the chunk, which is only expressed fully by the real-word conditions, starting with 2-word chunks, and which increases in complexity with chunk length). This design therefore gives rise to the eight hypothetical response profiles depicted in **Figure 2**. For example, a selectively syntactic region (–Lex, +Syn, – Sem) should respond identically across real-word and Jabberwocky conditions; a selectively combinatorial-semantic region (–Lex, –Syn, +Sem) should show a length effect (stronger responses to longer chunks) only in the real-word conditions; and a combined lexical, syntactic and combinatorial-semantic region (+Lex, +Syn, +Sem) should show length effects in both real-word and Jabberwocky conditions, with a stronger length effect in the real-word conditions. PDD’s design therefore permits empirical discrimination of different logically possible patterns of (in)sensitivity to lexical, syntactic, and combinatorial-semantic dimensions of language, with major implications for our understanding of the neural substrates that enable language comprehension.

**Figure 2.**
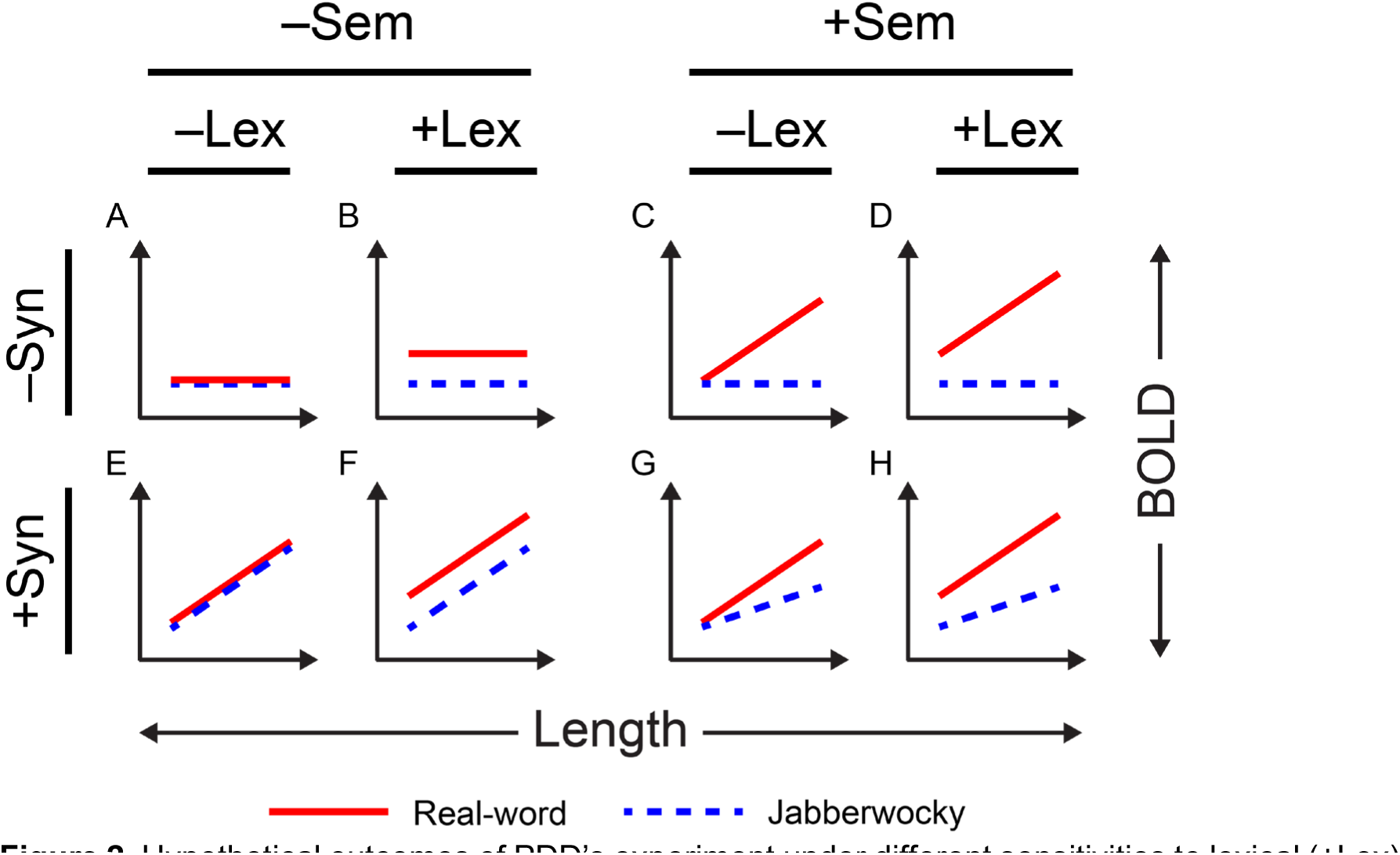
Hypothetical outcomes of PDD’s experiment under different sensitivities to lexical (±Lex), syntactic (±Syn), and combinatorial-semantic (±Sem) dimensions of language. Lexical processing (+Lex) predicts a larger overall response to real-word conditions than to Jabberwocky conditions, shifting all estimates for real words conditions upward. Syntactic processing (+Syn) predicts an increase in response to chunk length (*x*-axis) in both the real-word and Jabberwocky conditions. Combinatorial-semantic processing (+Sem) predicts a greater response to chunk length in the real-word conditions than the Jabberwocky conditions. These predictions combine to yield eight logically possible response profiles, many of which can be distinguished by testing for differences by condition type between the intercept (overall response) and/or slope (strength of response to chunk length).

PDD reported three key findings of relevance to the neural substrates of syntactic and semantic processing. **Finding 1**: Inferior frontal and posterior temporal language regions in the left hemisphere responded more strongly to longer constituents, even in the meaningless Jabberwocky conditions. **Finding 2**: The *slope* of this increase with chunk length was indistinguishable in these areas between the real-word and Jabberwocky conditions. The impact of this finding was plausibly enhanced by the additional apparent absence of a difference in *intercept* between conditions, such that the overall response profiles in these regions were nearly identical in the two types of conditions (similar to the selectively syntactic, –Lex, +Syn, –Sem, profile in **Figure 2**). **Finding 3**: By contrast, in anterior temporal and temporoparietal language regions, activation increased with chunk length in the real-word conditions but not the Jabberwocky conditions, with a significant difference in slope between the two condition types (similar to the –Syn, +Sem profiles in **Figure 2**). These findings have been reinforced by other studies showing syntactic/semantic dissociations with a similar topography to that reported by PDD (e.g., refs. (18, 19)).

In addition to their support for neurobiological effects of syntax in general (Finding 1), these findings have had a major influence on thinking about the division of labor within the human language system, which we group into two broad claims that were made directly by PDD or attributed to them by subsequent work. **Syntactic Hubs**: Finding 2 has been taken to support the existence of abstract syntactic hubs in inferior frontal and posterior temporal cortex (11, 13, 20–24). Because PDD reported qualitatively identical response profiles in these regions for real-word and Jabberwocky conditions, prior invocations of this empirical finding are often ambiguous between a strong form in which these hubs exclusively encode abstract combinatorics—with no reference to lexical or combinatorial-semantic content (refs. (11, 21, 25); profile –Lex, +Syn, –Sem in **Figure 2**)—and a weaker form in which these hubs do not encode combinatorial semantics, but may nonetheless respond more strongly to real words than pseudowords overall (ref. (18); profile +Lex, +Syn, –Sem in **Figure 2**). **Lexico-Semantic Hubs**: Finding 3 has been taken to support a selective role for anterior temporal and temporoparietal areas in lexical and combinatorial-semantic processing (refs. (5, 13, 16, 26–32); profile +Lex, –Syn, +Sem in **Figure 2**). For elaboration on the ways in which PDD’s study has influenced subsequent thinking about the neurobiology of language, see **SI Section 1**.

However, these claims now face empirical and methodological objections. Empirically, the existence of syntactic hubs (or, at least, the strong form of this claim) has been challenged by evidence of lexical processing in the inferior frontal and posterior temporal areas identified by PDD as abstract syntactic hubs (e.g., refs. (18, 33–37)), and the existence of lexico-semantic hubs has been challenged by evidence of sensitivity to structure in Jabberwocky materials in anterior temporal regions argued by PDD to be insensitive to such effects (e.g., refs. (33–35, 38, 39)). These prior studies raise concerns about the robustness and replicability of PDD’s reported pattern. Methodologically, some of the choices in PDD’s design and analyses are problematic. First, PDD used a between-subjects design to compare the real-word and Jabberwocky conditions (thus simultaneously varying both the sample of participants and the condition), even though this manipulation is feasible to perform in a within-subjects design that avoids this confound. Because individuals and, by extension, groups of individuals vary along numerous trait and state dimensions that are known to affect neural responses (e.g., refs. (40–42)), the magnitudes of neural responses in two groups cannot be confidently attributed to differences/similarities between conditions. Second, PDD used the same data both to define the regions of interest and to quantify their responses, introducing circularity (43). Third, PDD relied on traditional group analyses (40), which assume voxel-wise correspondence across individual brains. Ample evidence now exists for substantial inter-individual variability in the precise locations of functional areas in the association cortex (e.g., refs. (44–46)), including in the language network (e.g., refs. (33, 47)). Given that some of PDD’s claims rely on not finding certain effects in certain brain regions, the choice of traditional group analyses, which suffer from low sensitivity (48), is suboptimal. We stress that PDD’s approach and claims were reasonable for the time, and that some of the concerns above arise from empirical findings or methodological insights that were contemporaneous or subsequent to PDD’s publication date. However, because PDD’s findings continue to exert substantial influence, it is important to consider them in light of subsequent developments.

Motivated by these concerns, and in line with current emphasis in the field on robustness and replicability (49–54), we conduct three fMRI experiments (across a total of n=75 participants) that constitute the closest effort to date to replicate PDD’s original study while addressing the methodological issues above. First, we use a strictly within-subjects design. Second, we use independent data to define the regions of interest and to quantify their responses to the critical conditions. And third, we define regions of interest functionally in individual brains (e.g., refs. (33, 55, 56)), which has been shown to yield higher sensitivity and higher functional resolution (e.g., refs. (48, 57–59)).

We strongly replicate PDD’s key discovery of a basic chunk length effect in all experiments (see ref. (60) for another recent replication by another research group): activity in multiple language areas increases parametrically with the increasing length of linguistic context, even in the absence of lexical content. However, our results challenge the existence of both syntactic and semantic hubs. In particular, (a) all language regions with the exception of the language fROI in the TPJ / angular gyrus show a length effect in Jabberwocky conditions, (b) all language regions show an effect of ‘lexicality’, with real-word conditions eliciting stronger responses than Jabberwocky conditions, and (c) all language regions but the PostTemp language fROI show a length by lexicality interaction whereby the length effect is stronger in the real-word conditions compared to Jabberwocky conditions. We further show that these length effects do not critically depend on syntactic constituency per se but rather on the length of contiguous coherent text, which undermines PDD’s claim that syntactic constituency critically drives the length effect.

These findings challenge a bifurcation of the language system into discrete syntactic and lexico-semantic components. Our results instead join a growing body of evidence for an integrated network for language in the human brain (33, 59, 61, 62) within which internal specialization is primarily a matter of degree rather than kind (63–67), in contrast with influential proposals that posit a sharp separation between different types of linguistic representations and processes (3–5, 7).

## Results

We revisit PDD’s claims in three experiments. Experiment 1 seeks to replicate the finding of an overall increase in the BOLD response of language brain areas as a function of chunk length. Experiment 2 is a conceptual replication of PDD, including all of the original real-word conditions and a critical subset of the Jabberwocky conditions, as well as PDD’s two additional “non-constituent” conditions consisting of 3- and 4-word chunks that do not form valid syntactic constituents. Unlike PDD, in Experiment 2, and other experiments, we independently localize the language network in each participant and use a fully within-subjects design. Experiment 3 more directly targets the centrality of constituency for obtaining the length effect (stronger responses to longer chunks) by presenting participants with 24-word and 30-word stimuli that are composed of chunks of varying length (taken from naturalistic texts) which overwhelmingly do not form constituents in their source sentences (86.5% of the time; because Experiment 3 was originally designed with a different research goal in mind, avoiding constituents entirely was not a consideration). For details about these experiments, see **Materials & Methods**. Results are visualized in **Figure 3** (full significance testing details are given in **Table S1**), See **SI Section 9** for evidence that the results hold when we use the masks from PDD to define the language areas. See **SI Section 10** for evidence that the extremes of the length conditions—(jab-)c01 and (jab-)c12—replicate an established pattern of response in the language network. See **SI Section 11** for exploratory analyses of the right-hemisphere homotopes of the left-hemisphere language areas.

**Figure 3.**
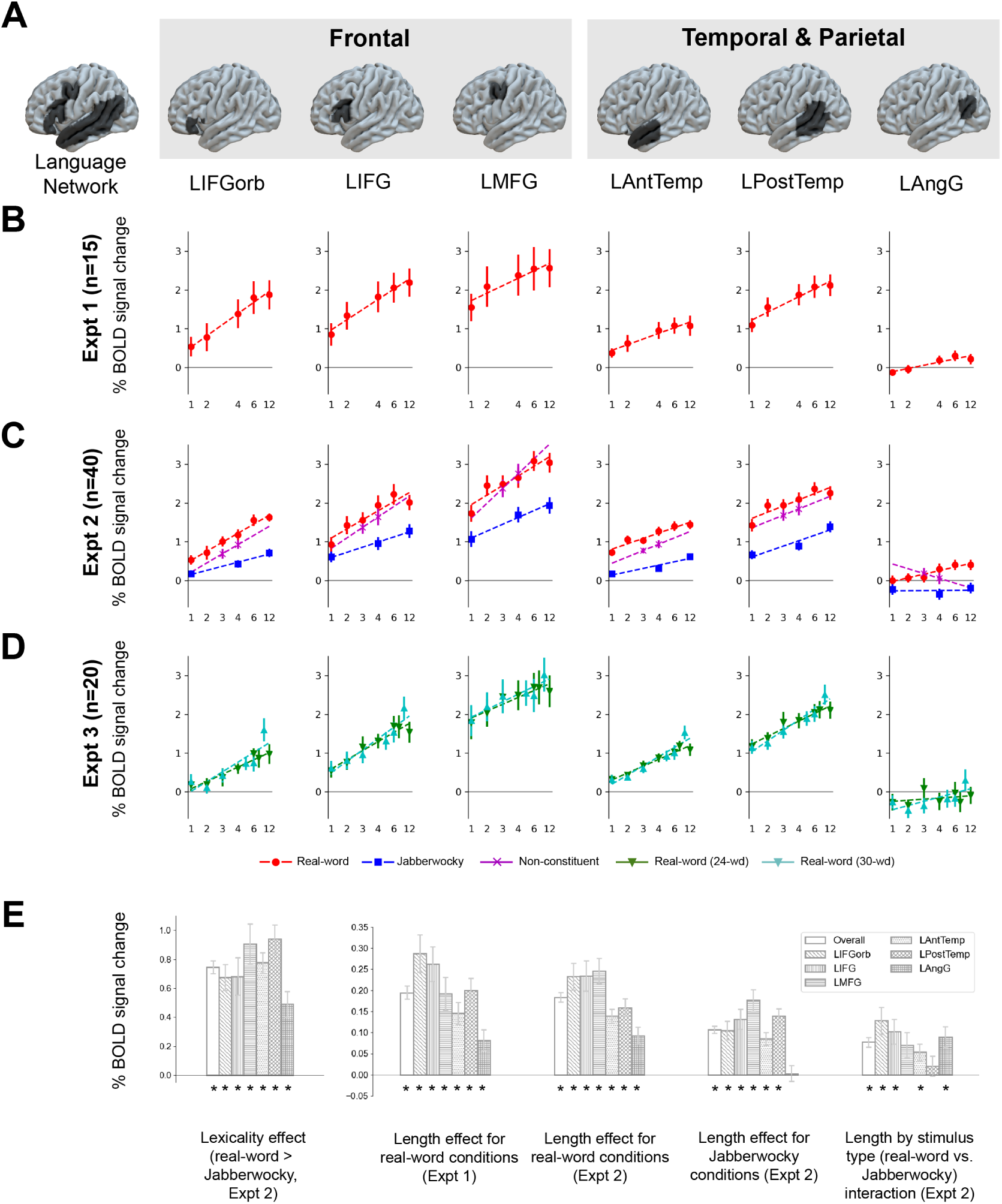
A. Group masks bounding the six left-hemisphere regions of the language network. The top 10% of language-responsive voxels (i.e., voxels that respond to the localizer contrast, sentences>nonwords) are selected within each mask in each participant (see Methods). **B.** Estimated response to each real-word condition in Expt 1 (which did not include Jabberwocky conditions). Responses in all regions increase with chunk length. **C.** Estimated response to each real-word, Jabberwocky, and non-constituent condition in Expt 2. Responses in all regions increase with chunk length in the real-word conditions, and responses in all regions but LAngG increase with chunk length in the Jabberwocky and non-constituent conditions. **D.** Estimated response to each condition of both the 24-word and 30-word items of Expt 3, both of which consisted of contiguous real-word chunks that generally did not form syntactic constituents. Responses in all regions increase with chunk length to a similar degree as in the real-word conditions of Expts 1 and 2. **E.** Key contrasts by language network fROI (left-to-right): overall lexicality effect (increase in response to real-word over Jabberwocky conditions in Expt 2, averaging over chunk length); length effect for real-word conditions in Expt 1 (slope of the line by participant from **B**); length effect for real-word conditions in Expt 2 (slope of the red line by participant from **C**); length effect for Jabberwocky conditions in Expt 2 (slope of the blue line by participant from **C**); increase in length effect in real-word conditions over Jabberwocky in Expt 2 (difference between the slopes of the red and blue lines by participant from **C**). Starred bars indicate statistically significant effects by likelihood ratio test (corrected for false discovery rate across fROIs; (136); see also **Supplementary Table S1**). Error bars show standard error of the mean over participants.

### Do the language regions show length effects?

For the real-word conditions, all regions show the pattern reported by PDD: significantly increasing activation as a function of chunk length, including a smaller increase at larger lengths (e.g., c06 to c12) in all three experiments (**Figure 3B, C, D, E**).

### Are there syntactic hubs that respond identically to the real-word and Jabberwocky conditions?

No language region shows the pattern (reported by PDD for inferior frontal and posterior temporal areas) of visually indistinguishable increases in neural activity with chunk length in the real-word and Jabberwocky conditions. Instead, all language regions’ responses are modulated by lexicality, either in the overall response, in the slope of the length effect, or both. Thus, no region appears to be a hub for abstract (i.e., content-independent) combinatorics.

### Do anterior temporal and temporoparietal language regions only show length effects in the real-word conditions?

We find a significant length effect in the Jabberwocky conditions for the language network as a whole, as well as for each region within it except for the temporoparietal LAngG region. Contrary to PDD’s claim that the anterior temporal language area (LAntTemp) is not responsive to chunk length in meaningless Jabberwocky materials, we find this effect robustly.

However, the LAngG region (which corresponds to PDD’s “TPJ” region; **Supplementary Figure S1**) only shows a length effect in the real-word conditions, as PDD claimed, and in direct pairwise comparisons between regions, the length effect for Jabberwocky stimuli is significantly weaker in the LAngG region than in all other language regions. Nonetheless, the length effect for real-word stimuli is also significantly weaker in LAngG than in all other language regions except for the LAntTemp region. Together with prior evidence (e.g., (59, 68–70)), this qualitative difference in response suggests that the LAngG region may not be part of the core language network (see Discussion).

### Are inferior frontal and posterior temporal language regions insensitive to combinatorial semantics, over and above syntax?

We find a significantly steeper slope for the length effect in the real-word conditions relative to the Jabberwocky conditions (length by stimulus type interaction) in the language network as a whole, as well as in each region within it except for the LMFG and LPostTemp regions. The length by stimulus type interaction in the LMFG region is positive and similar in magnitude to that of other regions, but it fails to reach significance. By contrast, the length by stimulus type interaction in the LPostTemp region is numerically near zero. This finding is contrary to PDD’s claim that the inferior frontal language areas (LIFGorb and LIFG) are equally sensitive to chunk length in real-word and Jabberwocky conditions (**Figure 3C, E**). However, the LPostTemp region shows highly similar length effects for real-word and Jabberwocky stimuli, as PDD claimed, and in direct comparisons, the difference in length effect between the real-word conditions and the Jabberwocky conditions is significantly weaker in the LPostTemp region relative to both the LIFGorb and the LIFG regions, the latter of which has been classically associated with syntactic processing (e.g., refs. (3, 71–73)). This result supports PDD’s claim that the LPostTemp region is equally sensitive to syntactic structure, with or without lexical content. We return to this finding in the **Discussion**.

### Does syntactic constituency critically drive the length effect?

The length effect in Experiment 2 is at least as strong in the non-constituent conditions as it is in the real-word constituent conditions, which undermines PDD’s claim that length effects are driven primarily by syntactic constituency. This finding is reinforced by Experiment 3, which evaluates length effects in materials composed primarily (86.5%) of non-constituents (**Figure 3D**). As shown, the length effect in response to these largely non-constituent materials is qualitatively similar to the length effects reported in Experiments 1 and 2, and quantitatively, we observe no significant differences in any region, or in the language network as a whole, between the length effect in Experiment 3 vs. in either Experiment 1 or Experiment 2 in between-group comparisons. Thus, syntactic constituency does not critically drive the length effects in the language network.

### Summary

Our results support a distributed burden of lexical, syntactic, and combinatorial-semantic processing throughout the language network (rather than the dissociation between syntactic and lexico-semantic sub-networks, as claimed by PDD), and challenge the claim that stronger responses to longer chunks are driven by syntactic constituency (given that these length effects are equally strong regardless of whether the chunks form constituents). The key similarities and differences between our findings and PDD’s are summarized in **Table 1**.

**Table 1.** Summary of key similarities and differences between PDD’s findings and those of our study with respect to sensitivity to lexical content, syntactic structure, and semantic composition. PDD reported (a) one set of regions (inferior frontal and posterior temporal) that were sensitive to structure (chunk length) in real-word conditions and equally sensitive to structure in Jabberwocky conditions (supporting abstract syntactic processing in these regions), and (b) another set of regions (anterior temporal and TPJ) that were sensitive to lexical content and insensitive to structure in Jabberwocky conditions. Our study does not reproduce several of PDD’s reported insensitivities (red minus signs) and challenges the purported double dissociation between lexical/semantic regions on the one hand (anterior temporal and temporoparietal areas) and abstract syntactic regions on the other (inferior frontal and posterior temporal areas). Instead, we find more broadly distributed lexical, syntactic, and combinatorial semantic effects throughout the language network, albeit with evidence (consistent with PDD’s claims) that the temporoparietal area is only sensitive to structure in real-word conditions and that the posterior temporal language area is equally sensitive to structure in both real-word and Jabberwocky conditions.

|  | Sensitivity to lexical content<br>(inconsistent with a purely syntactic function) |  | Sensitivity to structure in Jabberwocky<br>(inconsistent with an absence of syntactic function) |  | Greater sensitivity to structure in the presence of lexical content<br>(inconsistent with a selectively syntactic—vs. combinatorial semantic—function) |  |
| --- | --- | --- | --- | --- | --- | --- |
|  | PDD | This work | PDD | This work | PDD | This work |
| inferior frontal | — | + | + | + | — | + |
| anterior temporal | + | + | — | + | + | + |
| posterior temporal | — | + | + | + | — | — |
| AngG/TPJ | + | + | — | — | + | + |

## Discussion

Whether different brain areas specialize for different types of linguistic processing is a long-standing open question in the neurobiology of language. Perhaps the most frequently proposed pattern of specialization is a dissociation between some brain areas that selectively represent the structure of sentences and others that selectively represent the meaning (4, 5, 11, 74). This perspective is inspired in part by the well-known observation that some non-linguistic domains (e.g., mathematics, action planning, and music) also exhibit a kind of “syntax” in that they obey similar principles to language of sequential, hierarchical, and symbolic representation (9). If, as some have argued (8, 10, 75, 76), abstract syntactic composition is supported by a shared brain network with key loci in inferior frontal cortex, then the human capacity for language may derive from a more general capacity for structured symbol manipulation, which may in turn have arisen from anatomical changes to pre-frontal cortex during human evolution (11, 12). This position thus offers tantalizing continuities between language and other domains, along with explanatory links to evolutionary processes that might have set the stage for the emergence of language. However, the empirical literature that is used to support this position (from both neuroimaging and neuropsychology) widely assumes that spatial coordinates in the brain implement the same function across individuals—an assumption that is known to be incorrect for the language system and to lead systematically both to (i) failure to discover functional selectivities that are present in individual brains and (ii) conflation of functions that are distinct in individual brains (33, 55, 63, 77). This concern extends to the finding of distinct syntactic and lexico-semantic processing centers by PDD, whose results are additionally subject to concerns about (1) reliance on between-group comparisons to substantiate the claim of abstract syntactic processing and (2) using the same data to define the fROIs and to statistically examine their responses. Because PDD’s results have informed much subsequent theorizing about the neural basis of language and the structure of mental representations for language (e.g., refs. (11, 14–17)) and because of a growing effort in the field to replicate influential findings (49–54), here we revisit their claims across three fMRI experiments that address these methodological concerns.

Our findings robustly replicate PDD’s discovery of parametric sensitivity in language areas to the amount of linguistic context (increasing activation for longer spans of coherent text), as well as their finding that this pattern continues to hold in several areas even when lexical content is removed. Not only do we find this pattern across multiple experiments and in a different language (English) than the originally used French, but the effects are statistically indistinguishable across multiple independent groups of participants, which suggests that PDD uncovered a stable population-level signature of language comprehension in the brain (see also ref. (78)). This signature constitutes compelling evidence both that the brain’s response is modulated by linguistic complexity and that syntax contributes to this modulation independently of meaning. This finding from PDD (replicated here) is thus an important explanandum in any theory of the brain basis of language comprehension.

However, our findings do not accord with PDD’s proposed division of labor within the language network, namely, a double dissociation between syntactic and lexico-semantic sub-networks. Instead, our results reveal a more distributed pattern of lexical, syntactic, and combinatorial-semantic processing than that proposed by PDD (key similarities and differences between our findings and PDD’s are summarized in **Table 1**). First, our results challenge the notion of pure syntactic hubs (i.e., the claim that inferior frontal and posterior temporal language areas respond identically to syntactic complexity across real-word and Jabberwocky conditions). Instead, we find large and statistically significant increases in the language network’s response, including in the inferior frontal and posterior temporal areas, to real-word relative to Jabberwocky stimuli. This finding aligns with several prior studies (fMRI: ref. (33)—see **Figure S4** for a direct comparison of the overlapping subset of conditions, refs. (79, 80); intracranial recordings: ref. (78)) and with growing evidence for strong integration between structure and meaning in the representations and computations that underlie language processing across fields and approaches, from linguistic theory (e.g., refs. (81–84)), to psycholinguistics (e.g., refs. (85–88)), to computational linguistics (e.g., refs. (89–92)), to cognitive neuroscience (e.g., refs. (32, 35, 64, 78, 93–99)).

Second, our results challenge the claim that inferior frontal areas are insensitive to semantic (as opposed to purely syntactic) composition. Instead, we find larger increases in response to chunk length in the real-word compared to the Jabberwocky conditions in both the LIFG and LIFGorb language areas.

Third, our results challenge the claim that anterior temporal areas constitute lexico-semantic hubs that only process combinatorial structure in the presence of lexical meaning. Instead, we find significant increases in response to chunk length in the Jabberwocky conditions (see also refs. (33, 100); **Figure S1**).

Finally, our results challenge the centrality to the length effects of syntactic constituency (as opposed to other kinds of syntactic and semantic relations that hold between words in contiguous spans of language). Using PDD’s narrow contrast between 3- and 4-word chunks that do not form syntactic constituents, we find that the increase in brain activity from the 3-word condition to the 4-word condition is at least as large in the non-constituent stimuli as it is in the constituent stimuli in all regions except the LAngG language region (see below for discussion of this region). Furthermore, in a separate experiment that explored a wider range of implicit chunk lengths and consisted overwhelmingly (>86%) of non-constituent chunks, we find qualitatively and quantitatively similar effects of chunk length to those found when using valid syntactic constituents, with no significant difference in the length effect in any region or in the language network as a whole. This result is incompatible with PDD’s claim that chunk-length effects are driven primarily by the memory demands associated with assembling phrasal constituents, given that the same pattern of length effects arises from chunks that do not form constituents. Nonetheless, it is plausible that these length effects derive from linguistic complexity more broadly construed, and indeed we find that multiple independently motivated measures of predictability and memory demand during language processing correlate with chunk length (**SI Figure S2B**). Our results simply argue for an interpretation of length effects as driven by (perhaps diverse features of) richer linguistic *contexts*, rather than by phrasal constituency specifically. Other studies are needed to elucidate what those features are (see e.g., refs. (70, 100–108)).

An alternative conceptualization of these length effects draws on the framework of “proper” and “actual” domains of specialized information processing systems (109, 110), whereby the system’s degree of engagement with an input can be modulated by the degree of fit between a given input and the target domain for which the system is adapted. Given the highly combinatory and contextualized nature of natural language, several words of contiguous context may be necessary in order to identify a stimulus as “proper” to the language network. As a consequence, PDD’s shorter length conditions may fail by degrees to fully engage language processing mechanisms in the first place, thereby attenuating overall activation in the language system (see also (111)). Temporal receptive windows (TRWs, i.e., the length of the preceding context that affects the processing of the current input; refs. (112, 113)) could potentially serve as a filter for identifying domains proper to the language network, and indeed prior evidence supports the existence of TRWs for language on the order of a few words (24, 65, 78, 113, 114). However, the *causes* of these patterns of temporal receptivity are unknown, and they could derive from more basic kinds of linguistic processing (e.g., the degree to which nearby words can be composed into a syntactic parse may serve as a cue to whether an input is proper to the language network). In the absence of a deeper causal understanding of TRWs in the language network, viewing length effects as reflecting the distinction between proper and actual domains is not mutually incompatible with the interpretation whereby length effects reflect linguistic processing complexity.

Despite the lack of dissociation between syntactic and lexico-semantic processing centers and the broader distribution of diverse aspects of linguistic processing within the language network, our findings support two key functional asymmetries that were posited by PDD.

### 1) The LAngG/LTPJ language region differs functionally from the rest of the LH language network

First, like PDD, we find that the temporoparietal (LAngG in our terminology, LTPJ in PDD’s) language area behaves differently from the rest of the language regions: the length effect for Jabberwocky stimuli is (i) not significant and (ii) significantly smaller in the LAngG region than in all other language regions. Thus, the LAngG fROI is indeed less responsive to chunk length than other language regions in the absence of lexical content. We argue based on multiple converging lines of evidence that the difference between this region and the other language regions is because the LAngG/LTPJ language region is not in fact a core language area. Although this region shows robust responses to language (e.g., responding more to sentences than lists of pseudowords, refs. (13, 33)), it differs functionally from the core LH language network. *First*, the this region shows systematically weaker correlations with other language areas during naturalistic cognition paradigms than those areas do with each other (62, 68, 115). Furthermore, data-driven functional parcellation using dense individual-subject resting state data picks out the core temporal and frontal areas examined here—but not the LAngG/LTPJ region—as an integrated network, one that is highly overlapping with the one identified by task-based language localizers (59). *Second*, the LAngG/LTPJ region shows substantially weaker evidence than the other LH language areas of core language processing operations like next-word prediction and syntactic structure building (70, 105). *Third*, the LAngG/LTPJ region responds at least as strongly to pictures and videos of meaningful events as to sentences, and sometimes more strongly (69, 116). In addition, this region often shows *below-baseline* responses during language tasks (e.g., both in this study and in PDD), which could be because this area is instead a node in the *default mode network* (117–119), a brain network whose activity increases during rest, and which has been associated with high-level conceptual processing and episodic memory (117, 118, 120, 121). Many have argued that AngG broadly (cf. the language-responsive part of it) supports heteromodal conceptual integration (116, 122–129). This hypothesis could explain the greater response in the AngG/TPJ region to meaningful language stimuli, even in the absence of a selectively linguistic function.

### 2) The LPostTemp language region is sensitive to syntactic structure and word meanings but not combinatorial semantics

The claim from PDD that our results most strongly support is that the language-responsive area in the posterior temporal cortex is equally sensitive to structure, with or without lexical content. Although the overall response of the LPostTemp region to real-word stimuli is greater than its response to Jabberwocky stimuli, the difference in the length effect between real-word and Jabberwocky stimuli is virtually zero, as evidenced by similar slopes (**Figure 3C, E**)—the +Lex, +Synt, –Sem profile in **Figure 2**. This result is inconsistent with the strong characterization of LPostTemp as a pure syntactic hub, given that its response is strongly influenced by lexical content, independently of structure. However, it does suggest that the burden of combinatorial processing in the LPostTemp region is unaffected by the meaningfulness of the resulting structure, which supports a *lack of combinatorial-semantic processing over and above syntactic processing*. This profile appears to be unique to LPostTemp; the difference between the length effects in real-word vs. Jabberwocky stimuli is non-significant and near zero in LPostTemp, significant in inferior frontal language regions (LIFG and LIFGorb language regions), and significantly larger in the frontal regions than in LPostTemp in direct comparisons, even though LIFG is classically associated with abstract syntax (3, 71–73). This result is important for two reasons. First, it lends support to the hypothesis that the posterior temporal language area plays a special role in processing hierarchical syntax, relative to other language areas that frequently co-activate during language processing (5, 130). Second, it is to our knowledge the first clear evidence of region-level (cf. ref. (114)) functional differentiation within the human language network using localization methods that robustly account for inter-individual variation in the precise locations of language areas. These methods have so far yielded a highly distributed picture of linguistic (including sub-lexical, lexical, syntactic, and combinatorial-semantic) processing across the regions of the language network, with little evidence of network-internal structure (33, 70, 94, 105, 131, 132). Our current results support invariance in the LPostTemp region to combinatorial semantics (over and above syntax, –Sem in the terminology of **Figure 2**). However, invariance to combinatorial semantics is a weaker claim than the widespread interpretation of PDD as showing a selectively syntactic role for posterior temporal cortex in language processing (i.e., –Lex, –Sem, see **SI Section 1**).

In summary, contrary to PDD, we find lexicality effects in inferior frontal and posterior temporal language regions, length effects for Jabberwocky stimuli in the anterior temporal region, and length by lexicality interactions in the inferior frontal language regions. These results collectively challenge PDD’s hypothesized dissociation between language regions that selectively process abstract syntax and language regions that selectively process lexical and/or combinatorial semantics. Our results instead converge with growing evidence that linguistic representations and computations over a range of levels of description (phonological, lexical, syntactic, and combinatorial-semantic) are largely distributed across the language network (32, 35, 64, 65, 131, 133). We do find evidence of one key invariance claimed by PDD: although the posterior temporal language region is more responsive to materials with lexical content, it shows no increase in response to combinatorial semantics over and above syntax. This finding deserves further investigation, including with temporally-sensitive methods, to ask whether this brain region may support an earlier stage of comprehension that focuses on identifying the words and the grammatical relations among them, with inferences about e.g., logical semantics (entities, relations, quantifiers, entailments, etc.) taking place in other language areas. However, our results show that the burden of lexical, syntactic, and semantic processing is distributed across diverse cortical areas, and that no single area or set of areas constitutes the syntax hub claimed by PDD and related work.

## Materials and Methods

This study consists of three experiments. Experiment 1 focuses on the real-word conditions from PDD and attempts to replicate the basic length effect in the language network’s response. Experiment 2 additionally includes Jabberwocky conditions in order to test PDD’s critical theoretical claim: that a subset of the language network implements abstract, content-independent, syntactic processing. Experiment 3 targets the centrality of syntactic constituency by investigating length effects using chunks that overwhelmingly do not form syntactic constituents.

### Participants

Seventy-four unique individuals (age 18-38, 39 females) participated for payment (Experiment 1: n=15; Experiment 2: n=40, Experiment 3: n=20; one individual participated in both Experiment 2 and Experiment 3, on separate days). All but three subjects were right-handed—as determined by the Edinburgh Handedness Inventory (134), or self-report. All participants were native (age of acquisition <10 years old) or highly proficient (n=3) speakers of English (see ref. (62) for evidence that the language system of highly proficient speakers is similar to that of native speakers). All participants gave informed consent in accordance with the requirements of MIT’s Committee on the Use of Humans as Experimental Subjects (COUHES). Each participant completed a language localizer task (33) and a critical task.

### Critical Task

The design of Expts 1 and 2 followed PDD but used English materials available at https://osf.io/fduve/ (the original experiments were carried out in French). In particular, participants were presented with same-length stimuli (sequences of 12 words/nonwords), and the internal composition of these stimuli varied across conditions. The conditions in Experiment 1 were similar to PDD’s real-word conditions, except they did not include the 3-word constituent condition. Experiment 2 included three types of experimental manipulation that directly follow PDD’s original design: a) six real-word conditions: a sequence of twelve unconnected words (i.e., constituents of length 1: c01; here and elsewhere, our condition name abbreviations are similar to those in PDD), six 2-word constituents (c02), four 3-word constituents (c03), three 4-word constituents (c04), two 6-word constituents (c06), and a 12-word sentence (c12); b) three conditions that were a subset of the Jabberwocky conditions from PDD selected to span the range of constituent lengths: a list of twelve unconnected nonwords (jab-c01), three 4-word Jabberwocky constituents (jab-c04), and a 12-word Jabberwocky sentence (jab-c12); and c) two non-constituent conditions (four 3-word non-constituent chunks (nc03) and three 4-word non-constituent chunks (nc04)). Sample stimuli are shown in **Figure 1**.

Like the materials in Expts 1 and 2, the materials in Expt 3 implicitly contained sequences of contiguous chunks of varying length drawn from English sentences. However, unlike Expts 1 and 2, these chunks were not required to (and generally did not) form syntactic constituents in their source contexts. Thus, Expt 3 allows us to investigate the extent to which constituency is critical to the relationship between implicit chunk length and the brain’s response. The materials for Experiment 3 consisted of two sets, so as to span a large range of chunk lengths at a fine-grained level. Stimuli in set 1 were 24 words long in total and fell into length conditions based on the divisors of 24 (i.e., c01, c02, c03, c04, c06, c08, and c12. Stimuli in set 2 were 30 words long and fell into length conditions based on the divisors of 30 (i.e., c01, c02, c03, c05, c06, and c10).

### Procedure

The procedure was similar for the three experiments and followed PDD: participants saw the stimuli presented one word/nonword at a time in the center of the screen in all caps with no punctuation at the rate of 300 ms per word/nonword. In Experiment 1, the 150 trials (30 12-word stimuli x 5 conditions) were distributed across 5 runs, so that each run contained 6 trials per condition. In addition, each run included 108 s of fixation, for a total run duration of 216 s (3 min 36 s). In Experiment 2, the 330 trials (30 12-word stimuli x 11 conditions) were distributed across 10 runs, so that each run contained 3 trials per condition. In addition, each run included 121.2 s of fixation, for a total run duration of 240 s (4 min). In both experiments, the order of conditions and the distribution of fixation periods in each run were determined with the optseq2 algorithm (135). Experiment 3 used the same presentation format as Experiments 1 and 2, which means that the set 1 (24-word) trials lasted 7.2s, and set 2 (30-word) trials lasted 9s. The 156 trials of Expt 3 (12 24-word stimuli x 7 conditions plus 12 30-word stimuli x 6 conditions) were distributed across 6 runs, with each run containing 26 trials (14 24w trials, and 12 30w trials), 2 trials of each of the 13 conditions. Fixation periods were distributed as follows: 8 s at the beginning of the run, 5.4 s after each trial, and 8.2 s at the end of the run. Condition order varied across runs and participants, with the constraint that trials of the same condition did not appear in a row.

### Imaging, Functional Localization, and Data Analysis

Imaging, functional localization, and data analysis procedures are described in **SI Sections 2-6**.

## Author Contributions

Conceptualization: CS, HK, FM, EF; Design and materials creation: MS, HK, FM and EF; Experimental script creation: MS; fMRI data collection: HK, JA, MS, EF; fMRI data preprocessing and analysis: all authors; Formal statistical analysis: CS; Figures: CS; Writing original draft: CS and EF; Editing and comments: HK, CC, BL, and FM; Overall supervision: FM and EF.

## Classification

Biological Sciences – Neuroscience ; Social Sciences – Cognitive Sciences

## Acknowledgments

We acknowledge the Athinoula A. Martinos Imaging Center at the McGovern Institute for Brain Research, MIT. For technical support during scanning, we thank Steve Shannon and Atsushi Takahashi. We thank Tamar Regev for help with the ROI projection figures, the audience at the Neurobiology of Language conference in 2020 for helpful discussions of this work, and Rebecca Saxe, Ted Gibson, Nancy Kanwisher, and Stan Dehaene for comments on an earlier version of the manuscript. We also thank Christophe Pallier for sharing the ROI files. E.F. was supported by NIH awards R01-DC016607, R01-DC016950, and U01-NS121471, and by the funds from the McGovern Institute for Brain Research, Brain and Cognitive Sciences Department, and the Simons Center for the Social Brain. Conflicts of Interest: None.

## References

1. N. Chomsky, Aspects of the Theory of Syntax (MIT {P}ress, 1965).

2. L. Frazier, “Sentence Processing: A tutorial review” in Attention and Performance 12: The Psychology of Reading, M. Coltheart, Ed. (Erlbaum, 1987), pp. 559–586.

3. P. Hagoort, On Broca, brain, and binding: a new framework. Trends Cogn. Sci. 9, 416–423 (2005).

4. A. D. Friederici, Language in our brain: The origins of a uniquely human capacity (MIT Press, 2017).

5. I. Bornkessel-Schlesewsky, M. Schlesewsky, Reconciling time, space and function: a new dorsal--ventral stream model of sentence comprehension. Brain Lang. 125, 60–76 (2013).

6. H. Duffau, S. Moritz-Gasser, E. Mandonnet, A re-examination of neural basis of language processing: Proposal of a dynamic hodotopical model from data provided by brain stimulation mapping during picture naming. Brain Lang. 131, 1–10 (2014).

7. G. Hickok, D. Poeppel, The cortical organization of speech processing. Nat. Rev. Neurosci. 8, 393–402 (2007).

8. A. D. Patel, Language, music, syntax and the brain. Nat. Neurosci. 6, 674–681 (2003).

9. K. S. Lashley, The problem of serial order in behavior (Bobbs-Merrill Oxford, 1951).

10. E. Koechlin, T. Jubault, Broca’s Area and the Hierarchical Organization of Human Behavior. Neuron 50, 963–974 (2006).

11. S. Dehaene, F. Meyniel, C. Wacongne, L. Wang, C. Pallier, The neural representation of sequences: from transition probabilities to algebraic patterns and linguistic trees. Neuron 88, 2–19 (2015).

12. S. Dehaene, F. Al Roumi, Y. Lakretz, S. Planton, M. Sablé-Meyer, Symbols and mental programs: a hypothesis about human singularity. Trends Cogn. Sci. (2022).

13. C. Pallier, A.-D. Devauchelle, S. Dehaene, Cortical representation of the constituent structure of sentences. Proc. Natl. Acad. Sci. 108, 2522–2527 (2011).

14. J. J. Bolhuis, I. Tattersall, N. Chomsky, R. C. Berwick, How could language have evolved? PLoS Biol 12, e1001934 (2014).

15. W. T. Fitch, Toward a computational framework for cognitive biology: unifying approaches from cognitive neuroscience and comparative cognition. Phys. Life Rev. 11, 329–364 (2014).

16. I. Bornkessel-Schlesewsky, M. Schlesewsky, S. L. Small, J. P. Rauschecker, Neurobiological roots of language in primate audition: Common computational properties. Trends Cogn. Sci. 19, 142–150 (2015).

17. C. I. Petkov, E. Jarvis, Birds, primates, and spoken language origins: behavioral phenotypes and neurobiological substrates. Front. Evol. Neurosci. 4, 12 (2012).

18. W. Matchin, C. Hammerly, E. Lau, The role of the IFG and pSTS in syntactic prediction: Evidence from a parametric study of hierarchical structure in fMRI. Cortex 88, 106–123 (2017).

19. T. Goucha, A. D. Friederici, The language skeleton after dissecting meaning: A functional segregation within Broca’s area. Neuroimage 114, 294–302 (2015).

20. S. Dehaene, The Demodularization Hypothesis. The Neocortex 27, 293 (2019).

21. I. Hertrich, S. Dietrich, H. Ackermann, The role of the supplementary motor area for speech and language processing. Neurosci. Biobehav. Rev. 68, 602–610 (2016).

22. L. Wang, L. Uhrig, B. Jarraya, S. Dehaene, Representation of numerical and sequential patterns in macaque and human brains. Curr. Biol. 25, 1966–1974 (2015).

23. C. Pattamadilok, S. Dehaene, C. Pallier, A role for left inferior frontal and posterior superior temporal cortex in extracting a syntactic tree from a sentence. cortex 75, 44–55 (2016).

24. M. J. Nelson, et al., Neurophysiological dynamics of phrase-structure building during sentence processing. Proc. Natl. Acad. Sci. 114, E3669--E3678 (2017).

25. G. Kempen, Prolegomena to a neurocomputational architecture for human grammatical encoding and decoding. Neuroinformatics 12, 111–142 (2014).

26. K. Friston, G. Buzsáki, The functional anatomy of time: what and when in the brain. Trends Cogn. Sci. 20, 500–511 (2016).

27. M. A. Skeide, J. Brauer, A. D. Friederici, Brain functional and structural predictors of language performance. Cereb. Cortex 26, 2127–2139 (2016).

28. E. Zaccarella, A. D. Friederici, Merge in the human brain: A sub-region based functional investigation in the left pars opercularis. Front. Psychol. 6, 1818 (2015).

29. E. Zaccarella, M. Schell, A. D. Friederici, Reviewing the functional basis of the syntactic Merge mechanism for language: A coordinate-based activation likelihood estimation meta-analysis. Neurosci. Biobehav. Rev. 80, 646–656 (2017).

30. S. M. Wilson, et al., What role does the anterior temporal lobe play in sentence-level processing? Neural correlates of syntactic processing in semantic variant primary progressive aphasia. J. Cogn. Neurosci. 26, 970–985 (2014).

31. S. M. Frankland, J. D. Greene, Concepts and compositionality: in search of the brain’s language of thought. Annu. Rev. Psychol. 71, 273–303 (2020).

32. A. Bautista, S. M. Wilson, Neural responses to grammatically and lexically degraded speech. Lang. Cogn. Neurosci. 31, 567–574 (2016).

33. E. Fedorenko, P.-J. Hsieh, A. Nieto-Castañón, S. Whitfield-Gabrieli, N. Kanwisher, New method for fMRI investigations of language: defining ROIs functionally in individual subjects. J. Neurophysiol. 104, 1177–1194 (2010).

34. E. Fedorenko, A. Nieto-Castañón, N. Kanwisher, Syntactic processing in the human brain: What we know, what we don’t know, and a suggestion for how to proceed. Brain Lang. 120, 187–207 (2012).

35. E. Fedorenko, A. Nieto-Castañon, N. Kanwisher, Lexical and syntactic representations in the brain: An fMRI investigation with multi-voxel pattern analyses. Neuropsychologia 50, 499–513 (2012).

36. J. M. Rodd, M. H. Davis, I. S. Johnsrude, The neural mechanisms of speech comprehension: fMRI studies of semantic ambiguity. Cereb. Cortex 15, 1261–1269 (2005).

37. P. Hagoort, L. Hald, M. Bastiaansen, K. M. Petersson, Integration of word meaning and world knowledge in language comprehension. Science 304, 438–441 (2004).

38. C. Rogalsky, G. Hickok, Selective attention to semantic and syntactic features modulates sentence processing networks in anterior temporal cortex. Cereb. Cortex 19, 786–796 (2009).

39. C. Humphries, J. R. Binder, D. A. Medler, E. Liebenthal, Syntactic and semantic modulation of neural activity during auditory sentence comprehension. J. Cogn. Neurosci. 18, 665–679 (2006).

40. A. P. Holmes, K. J. Friston, Generalisability, Random Effects & Population Inference. Neuroimage 7, S754 (1998).

41. G. Chen, Z. S. Saad, J. C. Britton, D. S. Pine, R. W. Cox, Linear mixed-effects modeling approach to FMRI group analysis. Neuroimage 73, 176–190 (2013).

42. A. R. Hariri, The neurobiology of individual differences in complex behavioral traits. Annu. Rev. Neurosci. 32, 225–247 (2009).

43. N. Kriegeskorte, W. K. Simmons, P. S. F. Bellgowan, C. I. Baker, Circular analysis in systems neuroscience: The dangers of double dipping. Nat. Neurosci. 12, 535–540 (2009).

44. M. A. Frost, R. Goebel, Measuring structural--functional correspondence: spatial variability of specialised brain regions after macro-anatomical alignment. Neuroimage 59, 1369–1381 (2012).

45. A. M. Tahmasebi, et al., Is the link between anatomical structure and function equally strong at all cognitive levels of processing? Cereb. cortex 22, 1593–1603 (2012).

46. B. Vázquez-Rodríguez, et al., Gradients of structure--function tethering across neocortex. Proc. Natl. Acad. Sci. 116, 21219–21227 (2019).

47. K. Mahowald, E. Fedorenko, Reliable individual-level neural markers of high-level language processing: A necessary precursor for relating neural variability to behavioral and genetic variability. Neuroimage 139, 74–93 (2016).

48. A. Nieto-Castañón, E. Fedorenko, Subject-specific functional localizers increase sensitivity and functional resolution of multi-subject analyses. Neuroimage 63, 1646–1669 (2012).

49. H. Pashler, E.--J. Wagenmakers, Editors’ introduction to the special section on replicability in psychological science: A crisis of confidence? Perspect. Psychol. Sci. 7, 528–530 (2012).

50. R. O. Gilmore, M. T. Diaz, B. A. Wyble, T. Yarkoni, Progress toward openness, transparency, and reproducibility in cognitive neuroscience. Ann. N. Y. Acad. Sci. 1396, 5– 18 (2017).

51. M. C. Makel, J. A. Plucker, Facts are more important than novelty: Replication in the education sciences. Educ. Res. 43, 304–316 (2014).

52. Open Science Collaboration, Estimating the reproducibility of psychological science. Science (80-.). 349, aac4716 (2015).

53. D. J. Simons, The value of direct replication. Perspect. Psychol. Sci. 9, 76–80 (2014).

54. T. Yarkoni, J. Westfall, Choosing prediction over explanation in psychology: Lessons from machine learning. Perspect. Psychol. Sci. 12, 1100–1122 (2017).

55. R. Saxe, M. Brett, N. Kanwisher, Divide and conquer: a defense of functional localizers. Neuroimage 30, 1088–1096 (2006).

56. E. Fedorenko, The early origins and the growing popularity of the individual-subject analytic approach in human neuroscience. Curr. Opin. Behav. Sci. 40, 105–112 (2021).

57. E. Fedorenko, J. Duncan, N. Kanwisher, Language-selective and domain-general regions lie side by side within Broca’s area. Curr. Biol. 22, 2059–2062 (2012).

58. S. Shashidhara, F. S. Spronkers, Y. Erez, Individual-subject functional localization increases Univariate activation but not multivariate pattern discriminability in the “multiple-demand” frontoparietal network. J. Cogn. Neurosci. 32, 1348–1368 (2020).

59. R. M. Braga, L. M. DiNicola, H. C. Becker, R. L. Buckner, Situating the left-lateralized language network in the broader organization of multiple specialized large-scale distributed networks. J. Neurophysiol. 124, 1415–1448 (2020).

60. L. Giglio, M. Ostarek, K. Weber, P. Hagoort, Commonalities and asymmetries in the neurobiological infrastructure for language production and comprehension. Cereb. Cortex. Adv. online Publ. (2021).

61. T. L. Scott, J. Gallée, E. Fedorenko, A new fun and robust version of an fMRI localizer for the frontotemporal language system. Cogn. Neurosci. 8, 167–176 (2017).

62. S. Malik-Moraleda, et al., The universal language network: A cross-linguistic investigation spanning 45 languages and 11 language families. bioRxiv (2022).

63. E. Fedorenko, M. K. Behr, N. Kanwisher, Functional specificity for high-level linguistic processing in the human brain. Proc. Natl. Acad. Sci. 108, 16428–16433 (2011).

64. E. Fedorenko, I. Blank, M. Siegelman, Z. Mineroff, Lack of selectivity for syntax relative to word meanings throughout the language network. Cognition 203, 104348 (2020).

65. I. Blank, E. Fedorenko, No evidence for differences among language regions in their temporal receptive windows. Neuroimage, 116925 (2020).

66. E. Fedorenko, I. Blank, Broca’s Area Is Not a Natural Kind. Trends Cogn. Sci. (2020).

67. T. A. Keller, P. A. Carpenter, M. A. Just, The neural bases of sentence comprehension: a fMRI examination of syntactic and lexical processing. Cereb. cortex 11, 223–237 (2001).

68. I. Blank, N. Kanwisher, E. Fedorenko, A functional dissociation between language and multiple-demand systems revealed in patterns of BOLD signal fluctuations. J. Neurophysiol. 112, 1105–1118 (2014).

69. C. Shain, A. Paunov, X. Chen, B. Lipkin, E. Fedorenko, No evidence of theory of mind reasoning in the human language network. bioRxiv (2022).

70. C. Shain, I. A. Blank, E. Fedorenko, E. Gibson, W. Schuler, Robust effects of working memory demand during naturalistic language comprehension in language-selective cortex. J. Neurosci. (2022) https://doi.org/10.1523/JNEUROSCI.1894-21.2022.

71. A. Caramazza, E. B. Zurif, Dissociation of algorithmic and heuristic processes in language comprehension: Evidence from aphasia. Brain Lang. 3, 572–582 (1976).

72. A. D. Friederici, The brain basis of language processing: from structure to function. Physiol. Rev. 91, 1357–1392 (2011).

73. Y. Grodzinsky, P. Pieperhoff, C. Thompson, Stable brain loci for the processing of complex syntax: A review of the current neuroimaging evidence. cortex 142, 252–271 (2021).

74. G. Baggio, P. Hagoort, The balance between memory and unification in semantics: A dynamic account of the N400. Lang. Cogn. Process. 26, 1338–1367 (2011).

75. W. T. Fitch, M. D. Martins, Hierarchical processing in music, language, and action: Lashley revisited. Ann. N. Y. Acad. Sci. 1316, 87–104 (2014).

76. J. M. Novick, J. C. Trueswell, S. L. Thompson-Schill, Cognitive control and parsing: Reexamining the role of Broca’s area in sentence comprehension. Cogn. Affect. {\textbackslash}& Behav. Neurosci. 5, 263–281 (2005).

77. E. Fedorenko, N. Kanwisher, Neuroimaging of language: why hasn’t a clearer picture emerged? Lang. Linguist. Compass 3, 839–865 (2009).

78. E. Fedorenko, et al., Neural correlate of the construction of sentence meaning. Proc. Natl. Acad. Sci. 113, E6256--E6262 (2016).

79. M. Bedny, A. Pascual-Leone, D. Dodell-Feder, E. Fedorenko, R. Saxe, Language processing in the occipital cortex of congenitally blind adults. Proc. Natl. Acad. Sci. 108, 4429–4434 (2011).

80. W. Matchin, C. Brodbeck, C. Hammerly, E. Lau, The temporal dynamics of structure and content in sentence comprehension: Evidence from fMRI-constrained MEG. Hum. Brain Mapp. 40, 663–678 (2018).

81. R. Kaplan, J. Bresnan, “Lexical Functional Grammar: A Formal System for Grammatical Representation” in The Mental Representation of Grammatical Relations, J. Bresnan, Ed. (MIT Press, 1982), pp. 173–281.

82. C. Pollard, I. Sag, Head-driven Phrase Structure Grammar (University of Chicago Press, 1994).

83. R. Jackendoff, Semantic structures (The MIT Press, 1990).

84. A. Goldberg, Constructions at Work: the nature of generalization in language (Oxford University Press, 2006).

85. W. Schuler, A. Wheeler, Cognitive Compositional Semantics using Continuation Dependencies in Third Joint Conference on Lexical and Computational Semantics (*{{SEM}}’14), (2014).

86. Y. Kamide, C. Scheepers, G. T. M. Altmann, Integration of syntactic and semantic information in predictive processing: Cross-linguistic evidence from German and English. J. Psycholinguist. Res. 32, 37–55 (2003).

87. L. Pylkkänen, B. McElree, “The syntax-semantic interface: On-line composition of sentence meaning” in Handbook of Psycholinguistics, M. J. Traxler, M. A. Gernsbacher, Eds. (Elsevier, 2006).

88. M. C. MacDonald, N. J. Pearlmutter, M. S. Seidenberg, The lexical nature of syntactic ambiguity resolution. Psychol. Rev. 101, 676–703 (1994).

89. B.-D. Oh, W. Schuler, Contributions of Propositional Content and Syntactic Category Information in Sentence Processing in *Proceedings of the Workshop on Cognitive Modeling and Computational Linguistics*, (2021), pp. 241–250.

90. T. Mikolov, K. Chen, G. Corrado, J. Dean, Efficient Estimation of Word Representations in Vector Space. CoRR abs**/**1301.3, 1–12 (2013).

91. C. Dyer, A. Kuncoro, M. Ballesteros, N. A. Smith, Recurrent neural network grammars in Knight K, Lopez A, Mitchell M, Editors. Human Language Technologies. 2016 Conference of the North American Chapter of the Association for Computational Linguistics; 2016 June 12-17; San Diego (CA, USA).[Sl]: Association for Computational Linguistics (ACL), (2016).

92. C. Manning, H. Schütze, Foundations of Statistical Natural Language Processing (MIT Press, 1999).

93. C. Kauf, G. Tuckute, R. Levy, J. Andreas, E. Fedorenko, Lexical semantic content, not syntactic structure, is the main contributor to ANN-brain similarity of fMRI responses in the language network. bioRxiv, 2005–2023 (2023).

94. I. Blank, Z. Balewski, K. Mahowald, E. Fedorenko, Syntactic processing is distributed across the language system. Neuroimage 127, 307–323 (2016).

95. F. Keller, S. Gunasekharan, N. Mayo, M. Corley, Timing accuracy of web experiments: A case study using the {WebExp} software package. Behav. Res. Methods 41, 1 (2009).

96. A. J. Anderson, et al., Deep artificial neural networks reveal a distributed cortical network encoding propositional sentence-level meaning. J. Neurosci. 41, 4100–4119 (2021).

97. C. Caucheteux, A. Gramfort, J.-R. King, Disentangling syntax and semantics in the brain with deep networks in *International Conference on Machine Learning*, (2021), pp. 1336– 1348.

98. A. J. Reddy, L. Wehbe, Can fMRI reveal the representation of syntactic structure in the brain? Adv. Neural Inf. Process. Syst. 34, 9843–9856 (2021).

99. G. Merlin, M. Toneva, Language models and brain alignment: beyond word-level semantics and prediction. arXiv Prepr. arXiv2212.00596 (2022).

100. J. Brennan, et al., Syntactic structure building in the anterior temporal lobe during natural story listening. Brain Lang. 120, 163–173 (2012).

101. J. M. Henderson, W. Choi, S. G. Luke, R. H. Desai, Neural correlates of fixation duration in natural reading: evidence from fixation-related fMRI. Neuroimage 119, 390–397 (2015).

102. R. M. Willems, S. L. Frank, A. D. Nijhof, P. Hagoort, A. den Bosch, Prediction during natural language comprehension. Cereb. Cortex 26, 2506–2516 (2015).

103. J. Brennan, E. P. Stabler, S. E. Van Wagenen, W.-M. Luh, J. T. Hale, Abstract linguistic structure correlates with temporal activity during naturalistic comprehension. Brain Lang. 157, 81–94 (2016).

104. J. Brennan, J. T. Hale, Hierarchical structure guides rapid linguistic predictions during naturalistic listening. PLoS One 14, e0207741 (2019).

105. C. Shain, I. Blank, M. van Schijndel, W. Schuler, E. Fedorenko, fMRI reveals language-specific predictive coding during naturalistic sentence comprehension. Neuropsychologia 138, 107307 (2020).

106. A. Lopopolo, A. van den Bosch, K.-M. Petersson, R. M. Willems, Distinguishing syntactic operations in the brain: Dependency and phrase-structure parsing. Neurobiol. Lang., 1–64 (2020).

107. A. Lopopolo, S. L. Frank, A. den Bosch, R. M. Willems, Using stochastic language models (SLM) to map lexical, syntactic, and phonological information processing in the brain. PLoS One 12, e0177794 (2017).

108. M. Heilbron, K. Armeni, J.-M. Schoffelen, P. Hagoort, F. P. de Lange, A hierarchy of linguistic predictions during natural language comprehension. Proc. Natl. Acad. Sci. 119, e2201968119 (2022).

109. D. Sperber, The modularity of thought and the epidemiology of representations. Mapp. mind Domain Specif. Cogn. Cult., 39–67 (1994).

110. H. C. Barrett, R. Kurzban, Modularity in cognition: framing the debate. Psychol. Rev. 113, 628 (2006).

111. G. Tuckute, et al., Driving and suppressing the human language network using large language models. bioRxiv (2023).

112. U. Hasson, E. Yang, I. Vallines, D. J. Heeger, N. Rubin, A hierarchy of temporal receptive windows in human cortex. J. Neurosci. 28, 2539–2550 (2008).

113. Y. Lerner, C. J. Honey, L. J. Silbert, U. Hasson, Topographic Mapping of a Hierarchy of Temporal Receptive Windows Using a Narrated Story. J. Neurosci. 31, 2906–2915 (2011).

114. T. I. Regev, et al., Neural populations in the language network differ in the size of their temporal receptive windows. bioRxiv (2023) https://doi.org/10.1101/2022.12.30.522216.

115. A. Paunov, I. Blank, E. Fedorenko, Functionally distinct language and Theory of Mind networks are synchronized at rest and during language comprehension. J. Neurophysiol. 121, 1244–1265 (2019).

116. A. A. Ivanova, et al., The language network is recruited but not required for nonverbal event semantics. Neurobiol. Lang. 2, 176–201 (2021).

117. M. D. Greicius, B. Krasnow, A. L. Reiss, V. Menon, Functional connectivity in the resting brain: a network analysis of the default mode hypothesis. Proc. Natl. Acad. Sci. 100, 253– 258 (2003).

118. J. L. Vincent, et al., Coherent spontaneous activity identifies a hippocampal-parietal memory network. J. Neurophysiol. 96, 3517–3531 (2006).

119. M. E. Raichle, et al., A default mode of brain function. Proc. Natl. Acad. Sci. 98, 676–682 (2001).

120. J. Davey, et al., Exploring the role of the posterior middle temporal gyrus in semantic cognition: Integration of anterior temporal lobe with executive processes. Neuroimage 137, 165–177 (2016).

121. C. L. Philippi, D. Tranel, M. Duff, D. Rudrauf, Damage to the default mode network disrupts autobiographical memory retrieval. Soc. Cogn. Affect. Neurosci. 10, 318–326 (2015).

122. A. R. Price, M. F. Bonner, J. E. Peelle, M. Grossman, Converging evidence for the neuroanatomic basis of combinatorial semantics in the angular gyrus. J. Neurosci. 35, 3276–3284 (2015).

123. M. L. Seghier, The angular gyrus: multiple functions and multiple subdivisions. Neurosci. 19, 43–61 (2013).

124. M. F. Bonner, J. E. Peelle, P. A. Cook, M. Grossman, Heteromodal conceptual processing in the angular gyrus. Neuroimage 71, 175–186 (2013).

125. A. R. Price, J. E. Peelle, M. F. Bonner, M. Grossman, R. H. Hamilton, Causal evidence for a mechanism of semantic integration in the angular gyrus as revealed by high-definition transcranial direct current stimulation. J. Neurosci. 36, 3829–3838 (2016).

126. C. P. Davis, E. Yee, Features, labels, space, and time: Factors supporting taxonomic relationships in the anterior temporal lobe and thematic relationships in the angular gyrus. Lang. Cogn. Neurosci. 34, 1347–1357 (2019).

127. J. R. Binder, R. H. Desai, W. W. Graves, L. L. Conant, Where is the semantic system? A critical review and meta-analysis of 120 functional neuroimaging studies. Cereb. Cortex 19, 2767–2796 (2009).

128. E. Amit, C. Hoeflin, N. Hamzah, E. Fedorenko, An asymmetrical relationship between verbal and visual thinking: Converging evidence from behavior and fMRI. Neuroimage 152, 619– 627 (2017).

129. L. Fernandino, et al., Concept representation reflects multimodal abstraction: A framework for embodied semantics. Cereb. cortex 26, 2018–2034 (2016).

130. W. Matchin, G. Hickok, The cortical organization of syntax. Cereb. Cortex 30, 1481–1498 (2020).

131. T. I. Regev, et al., High-level language brain regions are sensitive to sub-lexical regularities. bioRxiv (2021).

132. F. Mollica, et al., Composition is the core driver of the language-selective network. Neurobiol. Lang. 1, 104–134 (2020).

133. C. Shain, I. A. Blank, E. Fedorenko, E. Gibson, W. Schuler, Robust effects of working memory demand during naturalistic language comprehension in language-selective cortex. J. Neurosci. 42, 7412–7430 (2022).

134. R. C. Oldfield, The assessment and analysis of handedness: the Edinburgh inventory. Neuropsychologia 9, 97–113 (1971).

135. A. M. Dale, B. Fischl, M. I. Sereno, Cortical surface-based analysis: I. Segmentation and surface reconstruction. Neuroimage 9, 179–194 (1999).

136. Y. Benjamini, D. Yekutieli, The control of the false discovery rate in multiple testing under dependency. Ann. Stat. 29, 1165–1188 (2001).

