## Supplementary Information for "Graded sensitivity to structure and meaning throughout the human language network"

### SI Section 1: Extended Discussion of Impact of Pallier, Devauchelle, & Dehaene (2011)

Here we provide an extended discussion of how the research community has tended to interpret Pallier, Devauchelle, & Dehaene (ref. (1); *PDD*) with respect to the neurobiological bases of syntactic vs. lexico-semantic processing.

PDD's finding of virtually identical parametric increases in inferior frontal and posterior temporal language areas' activation with chunk length across both real-word and Jabberwocky stimuli strongly suggests that these regions comprise an autonomous syntactic "module" (the –Lex, +Syn, –Sem profile in the terminology of **Figure 2** of the main article; ref. (2)). We believe this is the most straightforward interpretation of PDD's emphasis on "the **relative independence** of syntax from lexico-semantic features" (p. 2526, emphasis ours). Subsequent work by the authors has been more explicit about this interpretation: "Remarkably, when the stimuli were 'delexicalized' by substituting all content words with meaningless pseudowords while maintaining all grammatical words and inflections, a core set of areas in left IFG and pSTS **continued to respond identically**, suggesting their central role in the construction of abstract syntactic trees" (Dehaene et al., 2015, p. 12, emphasis ours; see also ref. (4)). This interpretation of PDD has been explicit in some studies (5) and is at least implied by other studies citing PDD in support of a "modular" (6), "core" (4, 7–10), or "pure" (11) syntax network. We therefore believe that an important component of PDD's influence has been the suggestion of an autonomous module or network for syntactic tree building that is insensitive to the *content* (meaning) of those trees, and that therefore responds identically to both real and Jabberwocky constituents.

This strong position is difficult to sustain in the face of abundant evidence that inferior frontal and posterior temporal language areas respond more to real-word than Jabberwocky stimuli (+Lex in **Figure 2**, e.g., refs. (12–16), *inter alia*). However, a weaker interpretation of PDD's inferior frontal and posterior temporal results focuses only on the absence of a difference in the *slope* of the parametric effect of constituent length between real-word and Jabberwocky stimuli, while allowing for a difference in overall response between the two condition types (the +Lex, +Syn, –Sem profile in **Figure 2**). This position abandons the notion that these regions constitute an encapsulated syntactic module (in PDD's words, the "independence of syntax from lexico-semantic features"), since they can be sensitive to the domain of application of syntactic processes—namely real vs. pseudoword constituents—and presumably are therefore involved in additional non-syntactic processes specific to real constituents, such as retrieving and representing lexical meanings. Under this view, the key invariance in these regions supported by PDD's results is to *combinatorial-semantic content*, given that the *increase* in activation with syntactic complexity is not greater in the real-word conditions (which have combinatorial semantic meaning) vs. the Jabberwocky conditions (which arguably do not). Studies that cite PDD in favor of syntax selectivity in these regions must at minimum have this weak interpretation in mind (e.g., refs. (6, 7, 11, 17–20)), although the distinction between the weak and strong claims above is rarely made explicit.

In addition, PDD's finding of a length effect in anterior temporal and temporoparietal language areas *only* in the real-word (but not the Jabberwocky) conditions has been taken to support a selectively semantic function for these areas (the +Lex, –Syn, +Sem profile in **Figure 2**), by PDD themselves and by work building on their results (21–29).

### SI Section 2: Data Acquisition, Preprocessing, and First-level Modeling

#### Data Acquisition

Whole-brain structural and functional data were collected using one of two configurations. Data from participants in Experiment 1 and Experiment 2 that were scanned prior to 2021 (n=40) were acquired on whole-body 3 Tesla Siemens Trio scanner with a 32-channel head coil at the Athinoula A. Martinos Imaging Center at the McGovern Institute for Brain Research at MIT. For these participants, T1-weighted, Magnetization Prepared Rapid Gradient Echo (MP-RAGE) structural images were collected in 176 sagittal slices with 1 mm isotropic voxels (TR = 2,530 ms, TE = 3.48 ms, TI = 900 ms, flip = 8 degrees). Functional, blood oxygenation level-dependent (BOLD) data were acquired using an EPI sequence with a 90° flip angle and using GRAPPA with an acceleration factor of 2; with the following parameters: thirty-three 4 mm thick near-axial slices acquired in an interleaved order (with 10% distance factor), with an in-plane resolution of 2.1 mm × 2.1 mm, FoV in the phase encoding (A >> P) direction 200 mm and matrix size 96 × 96, TR = 2000 ms and TE = 30 ms.

Data from participants in Experiment 2 and Experiment 3 that were scanned in 2021 or later (n=35) were collected on a whole-body 3 Tesla Siemens PRISMA scanner with a 32-channel head coil, also at the Athinoula A. Martinos Imaging Center at the McGovern Institute for Brain Research at MIT. For these participants, T1-weighted, MP-RAGE structural images were collected in 208 sagittal slices with 1 mm isotropic voxels (TR = 1,800 ms, TE = 2.37 ms, TI = 900 ms, flip = 8 degrees). Functional, BOLD data were acquired using an SMS EPI sequence with a 90° flip angle and using a slice acceleration factor of 2, with the following acquisition parameters: fifty-two 2 mm thick near-axial slices acquired in the interleaved order (with 10% distance factor), 2 mm × 2 mm in-plane resolution, FoV in the phase encoding (A >> P) direction 208 mm and matrix size 104 × 104, TR = 2,000 ms, TE = 30 ms, and partial Fourier of 7/8. For both functional sequences, the first 10 s of each run were excluded to allow for steady state magnetization.

#### Preprocessing

fMRI data were analyzed using SPM12 (release 7487), CONN EvLab module (release 19b) and other custom MATLAB scripts. Each participant's functional and structural data were converted from DICOM to NIFTI format. All functional scans were coregistered and resampled using B-spline interpolation to the first scan of the first session (30). Potential outlier scans were identified from the resulting subject-motion estimates as well as from BOLD signal indicators using default thresholds in CONN preprocessing pipeline (5 standard deviations above the mean in global BOLD signal change, or framewise displacement values above 0.9 mm, ref. (31)). Functional and structural data were independently normalized into a common space (the Montreal Neurological Institute [MNI] template; IXI549Space) using SPM12 unified segmentation and normalization procedure (32) with a reference functional image computed as the mean functional data after realignment across all timepoints omitting outlier scans. The output data were resampled to a common bounding box between MNI-space coordinates (-90, -126, -72) and (90, 90, 108), using 2mm isotropic voxels and 4th order spline interpolation for the functional data, and 1mm isotropic voxels and trilinear interpolation for the structural data. Last, the functional data were then smoothed spatially using spatial convolution with a 4 mm FWHM Gaussian kernel.

#### First-Level Modeling

For both the language localizer task and the critical task, effects were estimated using a General Linear Model (GLM) in which each experimental condition was modeled with a boxcar function convolved with the canonical hemodynamic response function (HRF) (fixation was modeled implicitly, such that all timepoints that did not correspond to one of the conditions were assumed to correspond to a fixation period). Temporal autocorrelations in the BOLD signal timeseries were accounted for by a combination of high-pass filtering with a 128 seconds cutoff, and whitening using an AR(0.2) model (first-order autoregressive model linearized around the coefficient a=0.2) to approximate the observed covariance of the functional data in the context of Restricted Maximum Likelihood estimation (ReML). In addition to main condition effects, other model parameters in the GLM design included first-order temporal derivatives for each condition (included to model variability in the HRF delays), as well as nuisance regressors controlling for the effect of slow linear drifts, subject-motion parameters, and potential outlier scans on the BOLD signal. Resulting effect estimates reflect percent BOLD signal change (PSC).

### SI Section 3: Participant-Specific Functional Localization

#### Procedure

Sixty-five participants (out of seventy-five total) performed the localizer task in the same session as the critical task, and the remaining participants performed the localizer in a different session; for evidence that localizer activations are stable across scanning sessions, see refs. (15, 33, 34). Most participants completed one or two additional tasks for unrelated studies. The entire scanning session lasted approximately two hours.

#### Localizer Task

The task used to localize the language network is described in detail in ref. (14). Briefly, we used a reading task that contrasted sentences and lists of unconnected, pronounceable nonwords in a standard blocked design with a counterbalanced order across runs. This contrast targets higher-level aspects of language including, critically, both lexico-semantic and syntactic/combinatorial processing, to the exclusion of perceptual (speech or reading-related) and articulatory processes (see e.g., ref. (35), for discussion). Stimuli were presented one word/nonword at a time. Participants were asked to read the materials attentively and to press a button at the end of each trial (included in order to help participants remain alert). Importantly, this localizer has been shown to generalize across different versions: the sentences > nonwords contrast, and similar contrasts between language and a degraded control condition, robustly activates the fronto-temporal language network regardless of the task, materials, modality of presentation, and particular language (14, 36–39). This includes generalization to both narrower contrasts (e.g., sentences > lists of unconnected words; refs. (14, 40)) and broader contrasts (e.g., listening to passages > listening to acoustically degraded passages (in fact, this localizer was used for two subjects in Experiment 3 due to unstable responses to the visual language localizer, as described below); e.g., refs. (36, 38)). Furthermore, the same network robustly emerges from naturalistic-cognition paradigms (e.g., resting state, listening to stories, watching movies) using the data-driven functional correlation approach (ref. (33), see also (41–43)), suggesting that this network constitutes a natural kind in the brain, and our localizer contrast is simply a quick and efficient way to identify this network as needed for testing critical hypotheses about it.

The whole-brain maps for the language localizer are available at: https://osf.io/fduve/.

#### Definition and Validation of Language-Responsive Functional Regions of Interest (fROIs)

For each participant (in each experiment), we defined a set of language-responsive functional ROIs (fROIs) using group-constrained, participant-specific localization (14). In particular, each individual participant's map for the sentences > nonwords contrast from the language localizer task was intersected with a set of six binary masks (the whole-brain maps for the language localizer are available at: https://osf.io/fduve/). These masks were derived from a probabilistic activation overlap map for the language localizer contrast in a large set of distinct participants (n=220) using the watershed parcellation, as described in ref. (14), and corresponded to relatively large areas within which most participants showed activity for the target contrast. These masks covered the fronto-temporal language network: three in the left frontal lobe falling within the IFG, its orbital portion, and the MFG, and three in the temporal and parietal cortex (**Figure 3A** of the main article). Within each mask, a participant-specific language fROI was defined as the top 10% of voxels with the highest *t*-values for the localizer contrast (see ref. (34) for evidence that the fROIs are similar when defined using a fixed statistical threshold).

For two participants, the data quality for the standard version of the language localizer was low; however, both had completed an alternative version of the localizer based on listening to short passages vs. acoustically degraded versions of those passages (see refs. (36, 38) for evidence that this version of the localizer identifies the same areas as the standard, reading-based localizer). For two additional participants, one run of the language localizer showed some fMRI artifacts; as a result, we used just one run for these participants (which is sufficient for identifying the language network).

Before examining the data from the critical experiments, we ensured that the language fROIs show the expected signature response (i.e., a stronger response to sentences than nonwords). To do so, we used an across-runs cross-validation procedure (e.g., ref. (45)), where one run of the localizer is used to define the fROIs, and the other run to estimate the responses, ensuring independence (e.g., ref. (46)). As expected, and replicating prior work (e.g., refs. (14, 15, 40, 47), *inter alia*), the language fROIs showed a robust sentences > nonwords effect across the 73 participants with cross-validated localizer contrast estimates (all $t_{(72)} > 7.26$, $p < 1e\text{-}9$, Cohen's $d > 0.085$), correcting for the number of regions (six) using the false discovery rate (FDR) correction (48). The localizer activation maps for the two additional participants with a single run of the localizer task (preventing across-runs cross validation) were evaluated by visual inspection and looked typical.

Our masks show a close correspondence with the group-level ROIs used in PDD (**Figure S1**, see **SI Section 9** for evidence that results replicate when using PDD's parcels as masks), with three exceptions: i) PDD did not recover the language-responsive region in the MFG (because this region consistently emerges in both contrast-based and functional-correlation-based analyses as part of the language network—e.g., refs. (14, 33, 41, 44)—we chose to include it here); ii) our AntTemp mask encompasses both of the anterior temporal ROIs in PDD (i.e., the anterior superior temporal sulcus (aSTS) ROI, and the temporal pole (TP) ROI) (because PDD found similar functional profiles for these two ROIs, and for ease of comparisons with past work from our group, we chose not to split our mask into two parts); and iii) our AngG mask only partially overlaps with PDD's TPJ (temporo-parietal junction) mask (however, none of PDD's critical claims that we challenge in the current manuscript pertain to this region; besides, our results are similar to PDD's in spite of this difference in the masks, see **SI Section 9**).

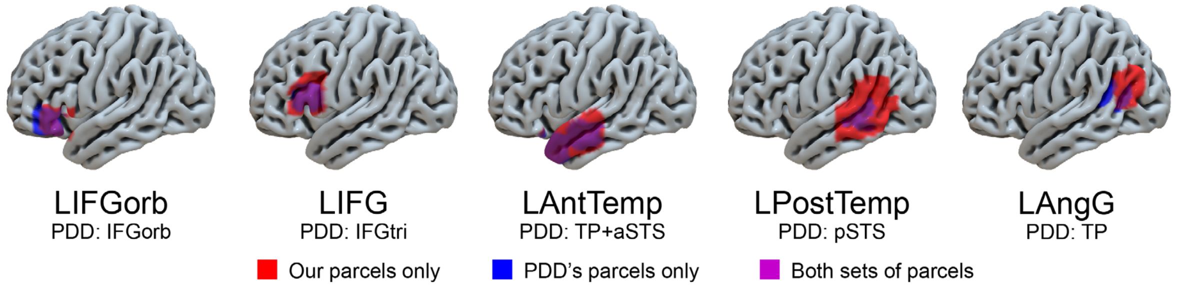

**Figure S1.** Visual comparison of language parcels used to define ROIs in PDD's original study vs. those used as anatomical masks for functional localization in our study. Red voxels are only included in our localizer masks (but not PDD's parcels), blue voxels are included in PDD's parcels (but not our localizer masks), and purple voxels are included both in PDD's parcels and in our localizer masks. Overlap between the two sets of parcels is generally high.

### SI Section 4: Materials Selection and Stimulus Design

**Experiments 1 and 2**: To create the materials for the real-word conditions, we extracted 180 2-word constituents (c02), 120 3-word constituents (c03), 90 4-word constituents (c04), 60 6-word constituents (c06), and 30 12-word constituents (c12) from the Penn-Treebank-parsed corpus (49) and the Natural Stories corpus (50). For each of the c02, c03, c04, and c06 conditions, the constituents were further manually concatenated into 30 12-word sequences, ensuring that syntactic or semantic dependencies would be unlikely to be formed across constituent boundaries. Finally, the c01 condition was created by selecting a set of 360 words from the full set of words in the Natural Stories corpus, and concatenating them into 30 12-word sequences, ensuring that adjacent words would be unlikely to combine syntactically or semantically.

To create the materials for the Jabberwocky conditions (jab-c01, jab-c04, and jab-c12), we took the strings from the c01, c04, and c12 real-word conditions and replaced all content words with pronounceable nonwords using the Wuggy software (51).

To construct the materials for the non-constituent conditions, we initially tried sampling 3- and 4-word non-constituent spans from the Natural Stories corpus (50), which contains hand-corrected phrase structure annotations. However, most strings extracted this way could often function as constituents in a different sentence context, especially given that many words in English can be used in multiple parts of speech. As a result, we hand-selected the non-constituent chunks from a larger set of texts and manually concatenated them to ensure that syntactic or semantic dependencies were unlikely to be formed across boundaries. We used an online book recommendation app (available at recommendmeabook.com) to sample the first page of classic and recent best-selling fiction books (e.g., 'The Poisonwood Bible' by Kingsolver). For every non-constituent chunk, we extracted a non-constituent of the appropriate length (3 or 4 words long, depending on the condition) from a book and then manually searched for another non-constituent that we believed would be unlikely to connect syntactically or semantically to the preceding one, and so on until the sequence (of four 3-word-long non-constituents, or three 4-word-long non-constituents) was complete. To protect against possible semantic dependencies, we often sampled non-constituents from different books for the same sequence. Using this method, we created 30 12-word sequences for each non-constituent condition (out of 120 3-word non-constituents for the nc03 condition, and out of 90 4-word non-constituents for the nc04 condition). Sample stimuli from Expts 1 and 2 are shown in **Figure 1** of the main article, and the full set of materials is available on OSF (https://osf.io/fduve/).

**Experiment 3**: To construct the materials for the (largely non-constituent) conditions of Experiment 3, we used the English Web Treebank of the Universal Dependencies (UD) corpus (52). First, we removed sentences that consisted of fewer than 17 words (to permit variability in the starting position of a chunk within the source sentence, even for the longest, 12-word, chunks), which resulted in a treebank of 6,273 sentences out of the original 16,622. These sentences were randomly assigned to conditions, and one chunk of the appropriate length was then extracted from each sentence, starting from a randomly chosen word index within the sentence from 1 to 5. We additionally required that: i) no token in a chunk could be a proper noun, a punctuation mark, a number (containing any digits), or a symbol; ii) no token in a chunk could have a miscellaneous/non-identified part-of-speech tag; iii) the first two characters of any word could not be capitalized (to avoid abbreviations); and iv) the chunk could not already be in the set of extracted chunks. We oversampled the number of sequences needed for each condition by a factor of three, to allow for subsequent filtering. We then filtered sequences *post hoc* to ensure that the sets of strings were matched across conditions (with a *p*-value of 0.05 or higher for any given condition pair in an independent-samples t-test) in terms of the following features: a) the average starting index of the string; b) the ratio of content to function words (where content words included nouns, verbs, adjectives, and adverbs); c) the average unigram lexical frequency; and d) the average word length (in letters). If any pair of conditions was not matched on one or more of these features, the 'worst offender' chunks were removed, and the statistics were recomputed. This was repeated until all pairs of conditions were matched on all features. The resulting set of chunks was then manually examined to remove chunks that straddled clausal boundaries or contained potentially sensitive, offensive, or highly culturally specific content. We then performed the matching described above more time on the set of approved chunks, to ensure that no biases were introduced by the content filtering.

Having selected the set of candidate chunks, we developed an algorithm to concatenate them into 24-word and 30-word items. Following PDD, the key desideratum was that the boundaries between adjacent strings within an item be clearly detectable. A long short-term memory language model was trained on the English Web Treebank from which the chunks were sampled. Using this model, for all chunks of a given length, each possible chunk pair combination was assigned a cost calculated as $log\left(\frac{p(s1+s2)}{p(s1)p(s2)}\right)$, where "s1 + s2" denotes chunk concatenation, and is computed by the language model. These costs were accumulated in an adjacency matrix. The chunk order with the minimum cost was found by greedily solving an asymmetric travelling salesman problem to select a minimum cost path through the chunks. This procedure resulted in a set of concatenated chunks to be used in the experiments. To create the condition with chunks of length 1, we used the words from the condition of chunk length 2 because the chunk length 1 condition should be most critically comparable to the next-length-up condition. However, because all conditions were well-matched for lexical properties, as described above, the words used in chunk length 1 condition were automatically matched to all the other conditions, too.

The full set of materials for Experiments 1-3 is available on OSF (https://osf.io/fduve/).

#### SI Section 5: Linguistic Features

We analyzed the materials in our real-words conditions in Experiments 1-2 with respect to six linguistic features with independent empirical support (*open nodes*, *node closings*, *storage cost*, *integration cost*, *5-gram surprisal*, and *PCFG surprisal*; all measures elaborated below), in order to shed light on possible causes of the length effects originally reported by PDD and replicated in our study. Results are reported in **Figure S2**. As shown, many of these features are either positively or negatively correlated with constituent length in these materials, suggesting potential directions for research that attempts to ground these effects in theory-driven accounts of language processing. We expand on these findings below.

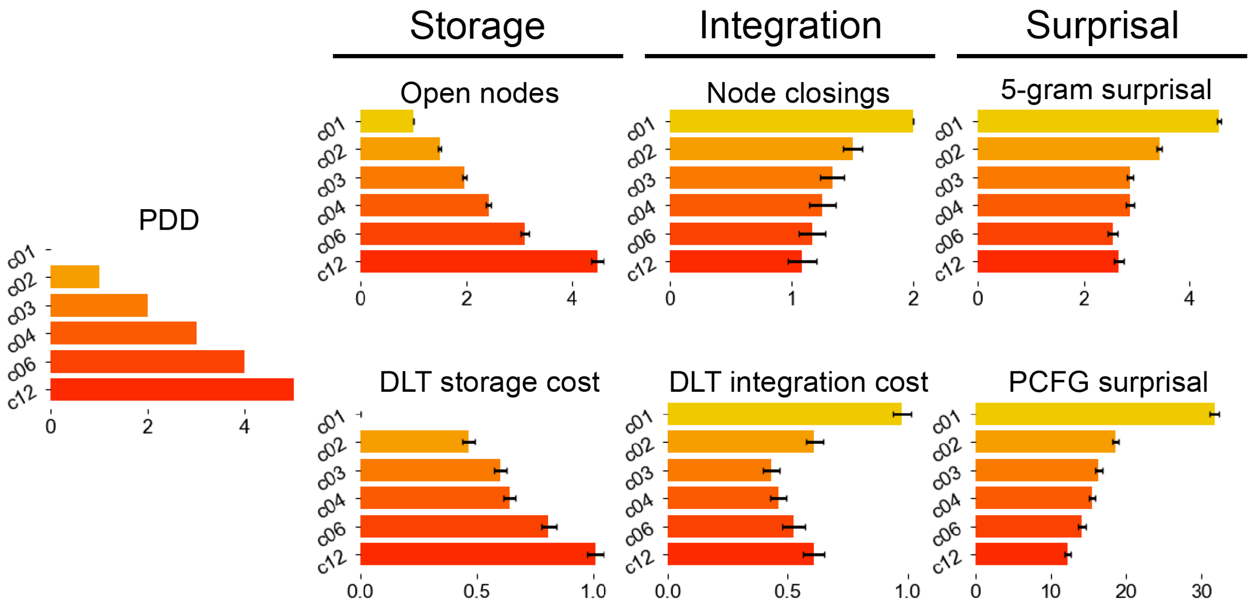

**Figure S2. B.** Mean value of linguistic features (memory- and surprisal-based) by constituent length for real-word conditions, compared to PDD-hypothesized monotonic increase (left). Error bars show standard errors of the mean across items.

##### Measures Derived from Memory-Based Accounts of Language Processing

**Open nodes** and **Node closings**: Ref. (7) elaborated on PDD's proposal—in the context of a follow-up study that used intracranial recordings—by hypothesizing a parsing mechanism that consumes more and more working memory until the constituent ends, permitting a Merge operation (53) whereby the memory allocated to representing that constituent is released and neural activation drops proportionately. Thus, PDD's pattern of stronger activity for sequences made up of longer constituents is hypothesized to derive from an accumulation of working memory demand by the parser over the course of constituent processing, with higher average demand for longer contiguous spans of text, since they can contain longer constituents. In ref. (7), these 'build-up' processes were encoded in the measure *open nodes* (a form of *storage cost* associated with maintaining items in working memory), and processes associated with merge and memory release were encoded in the measure *node closings* (a form of *integration cost* associated with retrieving and updating items in working memory).

We computed both of these measures as described in ref. (7) from phrase structure trees in a generalized categorial grammar (54) that were automatically generated for all stimuli using a probabilistic parser (55) and hand-corrected by an expert annotator (parses and annotations available from the ModelBlocks repository: https://github.com/modelblocks/modelblocks-release). For these and other measures, each distinct constituent within an item was treated as independent by the model. For sequences made up of multiple constituents, the values were averaged across constituents to derive a single value for each sequence.

Although *node closings* has independent psycholinguistic support (e.g., refs. (56–58)), this predictor is anticorrelated with PDD's constituent-length manipulations and therefore cannot explain the effect (**Figure S2, Node closings**). *Open nodes* is better correlated with PDD's expected pattern (**Figure S2, Open nodes**).

**Dependency Locality Theory (DLT) storage** and **integration cost:** The Dependency Locality Theory (DLT, ref. (59)) is one of many theories of working memory use in human sentence processing (see also e.g., refs. (60–62)). It was selected for analysis based on evidence that it best characterizes activity in the language network among a range of existing memory-based theories (63). DLT effects have also been reported in behavioral studies (64, 65). The DLT posits measures that are conceptually related to the *open nodes* (storage) and *node closings* (integration) predictors discussed above. In the DLT, *storage cost* (**Figure S2, Storage cost**) tracks the number of incomplete syntactic dependencies that must be maintained in memory. *Integration cost* (**Figure S2, Integration cost**) tracks the difficulty of constructing a syntactic dependency as a function of the number of intervening discourse referents. The measures of integration cost that we use here incorporate modifications described in ref. (66) that discount the cost of preceding modifiers and coordinate structures and increase the cost of verbs, following theoretical and empirical support described in ref. (63). DLT measures were computed automatically from the hand-corrected phrase structure trees described above.

The *storage cost* measure is correlated with PDD's constituent-length manipulation; the *integration cost* measure is anticorrelated with PDD's manipulation and therefore cannot explain the effect.

#### Measures Derived from Surprisal-Based Accounts of Language Processing

An alternative class of accounts of language comprehension with broad empirical support (e.g., refs. (67–71)) focus on the predictability of incoming words in context (72, 73). Here we focus on two such measures, following ref. (71), who found support for both in neural responses of the language system during naturalistic story comprehension.

**5-gram surprisal:** The negative log probability of a word in context as computed by KenLM 5-gram language models (74) from frequency counts in the Gigaword 3 corpus (75) (**Figure S2, 5-gram surprisal**). 5-gram models condition the probability distribution over the upcoming word on the sequence of 4 words that precede it, using default interpolation and backoff settings as described in ref. (74). 5-gram models capture local word co-occurrence statistics but struggle to capture effects of larger-scale syntactic structures (e.g., constituency, long-distance dependencies, etc.). Robust effects of 5-gram predictability (and related models) are consistently reported in both behavioral (67, 68, 76) and neuroimaging (69, 71, 77) studies.

**PCFG surprisal:** The negative log probability of a word in context as computed by the probabilistic context-free grammar (PCFG) parser of ref. (55), trained on trees from the Penn Treebank (49) that were automatically reannotated into a generalized categorial grammar (GCG) formalism (54) (**Figure S2, PCFG surprisal)**. PCFG models condition only on hypothesized syntactic analyses of sentences. They therefore excel at capturing syntactic influences on expectations but struggle to capture local word-to-word cooccurrence patterns. PCFG effects have been reported in both behavioral (78, 79) and neuroimaging (58, 71) studies.

Both surprisal measures are anticorrelated with PDD's constituent-length manipulation and therefore cannot explain the effect.

#### SI Section 6: Contrast Definition for the Critical Experiments

The first-level models estimate the response in PSC to each condition of the critical experiment (e.g., c02, jab-c12, etc.). However, our critical research questions aggregate over these conditions in different ways (Is the response to real-word stimuli bigger than the response to Jabberwocky stimuli overall? Does activity increase on chunk length? Etc.) Thus, as was done by PDD, we derive our key measures from the condition-level estimates. The resulting aggregate contrasts (estimated within each participant) are used as dependent variables for statistical analysis.

To estimate the overall response to real-word, Jabberwocky, and non-constituent conditions, we computed a by-participant average of the responses to the stimuli in each of these broader stimulus types. To estimate the difference in response between real-word and Jabberwocky conditions, we took the by-participant difference between the averages within those two stimulus types *only for lengths 1, 4, and 12*, which were represented for both stimulus types.

To estimate the parametric change in BOLD response as a function of constituent length, we computed the slope by participant of the best-fit line relating constituent length values to their associated first-level PSC estimates. To do so, we followed PDD in treating conditions c01, c02, c03, c04, c06, and c12 as equidistant, based on their observation of a sublinear monotonic relationship between length (in words) and the BOLD response. To model length effects in Expt 3, which includes conditions not present in PDD's original study (i.e., lengths 5, 8, and 10), we interpolated linearly between the points in PDD's original continuum. For example, length 5 (which was not used by PDD) was treated as lying halfway between lengths 4 and 6 (both of which were used by PDD). To estimate the difference in sensitivity to constituent length between stimulus types (e.g., between real-word and Jabberwocky conditions), we took the by-participant difference in slope between the two stimulus types.

#### SI Section 7: Statistical Analysis

We modeled the contrast values (as defined above, e.g., the by-participant difference in constituent length effect between real-word and Jabberwocky stimuli) as dependent variables in linear mixed effects models in `lme4` (80) when examining entire networks, with random effects for Participant and fROI, or simple linear models when examining the fROIs separately (since fROI-level models contain one contrast estimate per participant, there is no by-participant hierarchical structure to model). When examining the fROIs separately, reported *p*-values are adjusted for false discovery rate (48) over the number of fROIs in the network.

Network-wide contrast estimates were tested with the following mixed effects model:

```
Contrast ~ 1 + (1 | Participant) + (1 | fROI)
```

The critical variable in the above model is the intercept (**1**), which was tested by comparing this model (using a likelihood ratio test) to one in which the intercept is fixed at 0:

```
Contrast ~ 0 + (1 | Participant) + (1 | fROI)
```

Regional contrast estimates were tested (against zero) using an unpaired *t*-test. Pairwise tests of the difference in a contrast between two regions were tested in the same way, only using the *difference* in a given contrast from one region to the other (within an individual) as the dependent variable, rather than the contrast itself.

### SI Section 8: Full Statistical Results from the Main Article

| Contrast | Expt | fROI | β | σ(β) | t | *p* |
|---|---|---|---|---|---|---|
| Constituent length for real-word conditions | 1 | **Overall** | 0.19 | 0.04 | 5.34 | < 0.001*** |
| | 1 | **LIFGorb** | 0.29 | 0.04 | 6.51 | < 0.001*** |
| | 1 | **LIFG** | 0.26 | 0.04 | 6.39 | < 0.001*** |
| | 1 | **LMFG** | 0.19 | 0.04 | 4.91 | < 0.001*** |
| | 1 | **LAntTemp** | 0.15 | 0.03 | 5.62 | < 0.001*** |
| | 1 | **LPostTemp** | 0.20 | 0.03 | 6.80 | < 0.001*** |
| | 1 | **LAngG** | 0.08 | 0.03 | 3.28 | 0.013* |
| Constituent length for real-word conditions | 2 | **Overall** | 0.18 | 0.03 | 5.98 | < 0.001*** |
| | 2 | **LIFGorb** | 0.23 | 0.03 | 7.37 | < 0.001*** |
| | 2 | **LIFG** | 0.23 | 0.04 | 6.56 | < 0.001*** |
| | 2 | **LMFG** | 0.25 | 0.03 | 8.20 | < 0.001*** |
| | 2 | **LAntTemp** | 0.14 | 0.02 | 8.18 | < 0.001*** |
| | 2 | **LPostTemp** | 0.16 | 0.02 | 7.72 | < 0.001*** |
| | 2 | **LAngG** | 0.09 | 0.02 | 4.69 | < 0.001*** |
| Constituent length for Jabberwocky conditions | 2 | **Overall** | 0.11 | 0.03 | 4.08 | 0.003** |
| | 2 | **LIFGorb** | 0.11 | 0.03 | 3.97 | < 0.001*** |
| | 2 | **LIFG** | 0.10 | 0.02 | 4.78 | < 0.001*** |
| | 2 | **LMFG** | 0.13 | 0.02 | 5.48 | < 0.001*** |
| | 2 | **LAntTemp** | 0.18 | 0.03 | 6.97 | < 0.001*** |
| | 2 | **LPostTemp** | 0.09 | 0.02 | 5.58 | < 0.001*** |
| | 2 | LAngG | 0.14 | 0.02 | 8.30 | 1.000 |
| Lexicality effect (real-word > Jabberwocky) | 2 | **Overall** | 0.75 | 0.10 | 7.49 | < 0.001*** |
| | 2 | **LIFGorb** | 0.67 | 0.09 | 7.43 | < 0.001*** |
| | 2 | **LIFG** | 0.68 | 0.13 | 5.29 | < 0.001*** |
| | 2 | **LMFG** | 0.91 | 0.14 | 6.68 | < 0.001*** |
| | 2 | **LAntTemp** | 0.78 | 0.07 | 11.21 | < 0.001*** |
| | 2 | **LPostTemp** | 0.94 | 0.09 | 9.94 | < 0.001*** |
| | 2 | **LAngG** | 0.49 | 0.09 | 5.51 | < 0.001*** |
| Constituent-length by stimulus type (real-word vs. Jabberwocky) interaction | 2 | **Overall** | 0.08 | 0.02 | 3.28 | 0.004** |
| | 2 | **LIFGorb** | 0.13 | 0.03 | 4.03 | 0.004** |
| | 2 | **LIFG** | 0.10 | 0.03 | 3.47 | 0.006** |
| | 2 | LMFG | 0.07 | 0.03 | 2.31 | 0.078 |
| | 2 | **LAntTemp** | 0.05 | 0.02 | 2.63 | 0.045* |
| | 2 | LPostTemp | 0.02 | 0.02 | 0.84 | 1.000 |
| | 2 | **LAngG** | 0.09 | 0.02 | 3.68 | 0.005** |
| Length effect in Expt 3 (mostly non-constituents) | 3 | **Overall** | 0.19 | 0.03 | 5.72 | < 0.001*** |
| | 3 | **LIFGorb** | 0.21 | 0.03 | 6.98 | < 0.001*** |
| | 3 | **LIFG** | 0.26 | 0.04 | 6.82 | < 0.001*** |
| | 3 | **LMFG** | 0.18 | 0.03 | 5.59 | < 0.001*** |
| | 3 | **LAntTemp** | 0.20 | 0.02 | 9.87 | < 0.001*** |
| | 3 | **LPostTemp** | 0.23 | 0.03 | 9.04 | < 0.001*** |

| | | | | | | |
|---|---|---|---|---|---|---|
| | 3 | LAngG | 0.06 | 0.03 | 2.42 | 0.063 |
| Length effect in Expt 1 (constituents) vs. Expt 3 (mostly non-constituents) | 1 & 3 | Overall | 0.00 | 0.03 | -0.08 | 0.938 |
| | 1 & 3 | LIFGorb | -0.07 | 0.05 | -1.44 | 1.000 |
| | 1 & 3 | LIFG | 0.00 | 0.06 | -0.06 | 1.000 |
| | 1 & 3 | LMFG | -0.01 | 0.05 | -0.15 | 1.000 |
| | 1 & 3 | LAntTemp | 0.06 | 0.03 | 1.74 | 1.000 |
| | 1 & 3 | LPostTemp | 0.03 | 0.04 | 0.80 | 1.000 |
| | 1 & 3 | LAngG | -0.02 | 0.04 | -0.47 | 1.000 |
| Length effect in Expt 2 (constituents) vs. Expt 3 (mostly non-constituents) | 2 & 3 | Overall | 0.01 | 0.03 | 0.25 | 0.804 |
| | 2 & 3 | LIFGorb | -0.02 | 0.05 | -0.40 | 1.000 |
| | 2 & 3 | LIFG | 0.02 | 0.06 | 0.43 | 1.000 |
| | 2 & 3 | LMFG | -0.06 | 0.05 | -1.28 | 1.00 |
| | 2 & 3 | LAntTemp | 0.06 | 0.03 | 2.29 | 0.316 |
| | 2 & 3 | LPostTemp | 0.07 | 0.03 | 2.07 | 0.316 |
| | 2 & 3 | LAngG | -0.03 | 0.03 | -0.85 | 1.000 |
| Difference in constituent length effect for real-word conditions | 2 | **LAngG vs. LIFGorb** | 0.14 | 0.03 | 4.47 | < 0.001*** |
| | 2 | **LAngG vs. LIFG** | 0.14 | 0.04 | 3.87 | 0.002** |
| | 2 | **LAngG vs. LMFG** | 0.15 | 0.03 | 5.27 | < 0.001*** |
| | 2 | LAngG vs. LAntTemp | 0.05 | 0.02 | 1.91 | 0.146 |
| | 2 | **LAngG vs. LPostTemp** | 0.07 | 0.02 | 3.15 | 0.009** |
| Difference in constituent length effect for Jabberwocky conditions | 2 | **LAngG vs. LIFGorb** | 0.10 | 0.02 | 5.20 | < 0.001*** |
| | 2 | **LAngG vs. LIFG** | 0.13 | 0.02 | 5.60 | < 0.001*** |
| | 2 | **LAngG vs. LMFG** | 0.17 | 0.03 | 6.16 | < 0.001*** |
| | 2 | **LAngG vs. LAntTemp** | 0.08 | 0.02 | 4.76 | < 0.001*** |
| | 2 | **LAngG vs. LPostTemp** | 0.14 | 0.02 | 6.72 | < 0.001*** |
| Difference in constituent-length by stimulus type (real-word vs. Jabberwocky) interaction | 2 | **LPostTemp vs. LIFGorb** | 0.11 | 0.03 | 4.30 | 0.002** |
| | 2 | **LPostTemp vs. LIFG** | 0.08 | 0.03 | 3.27 | 0.021* |

**Table S1:** Significance tests of key contrasts with estimates, standard errors, and *t*-values (respectively columns β, σ(β), and t). *p*-values are generated by likelihood ratio tests of linear mixed-effects models, with by-fROI results corrected for false discovery rate (FDR) applied over all six fROIs using the Benjamini-Yekutieli procedure (48) using a nominal significance level of $\alpha = 0.05$. Starred *p*-values indicate statistical significance under FDR correction (*: p ≤ 0.05, **: p ≤ 0.01, *** p: ≤ 0.001). Significant regions are shown in **bold** in the fROI column.

### SI Section 9: Results Replicate When Using PDD's ROIs as Masks to Define the Language fROIs

This study constrained the participant-specific functional localization procedure using broad masks for language areas that have been validated in prior work (e.g., Fedorenko et al., 2010). As discussed in **SI Section 3**, five out of six of these masks correspond closely to the regions of interest (ROIs) reported in PDD, but the overlap between our masks and PDD's ROIs is not perfect. To ensure that our results are not due the choice of the particular masks, in this section we rerun our main analyses using PDD's language ROIs as anatomical localizer masks, rather than our standard anatomical masks. As shown in **Figure S3** and **Table S2**, results using PDD's parcels as localizer masks are highly similar to those reported using our standard masks in the main article, which indicates that results do not hinge critically on our choice of anatomical masks.

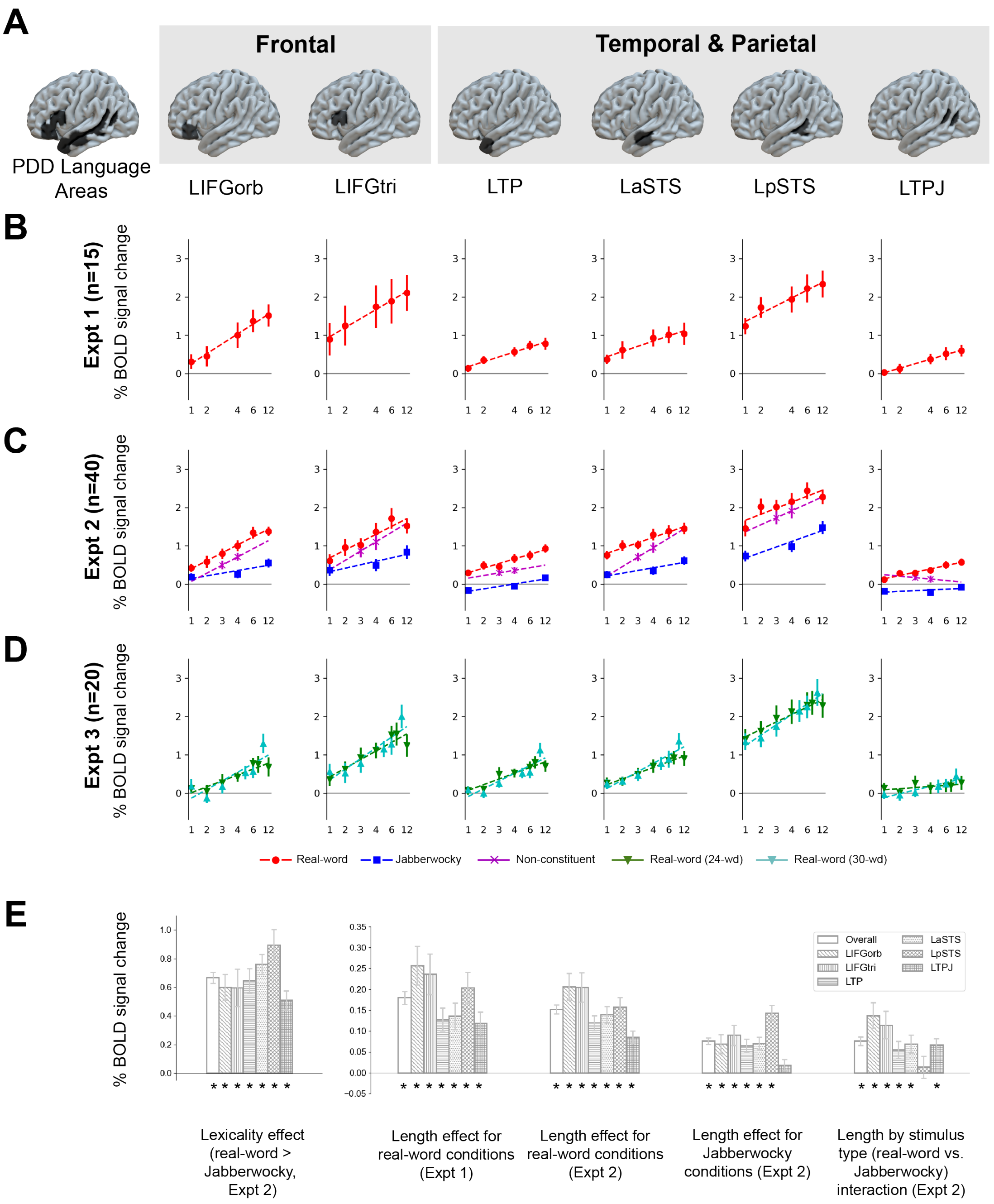

**Figure S3.** Main results (parallel of **Figure 3** of the main article) using PDD's anatomical ROIs as localizer masks, rather than the standard localizer masks. **A.** PDD's anatomical ROI parcels, used here as localizer masks. The top 10% of language-selective voxels are selected within each mask in each participant. **B.** Estimated response to each condition of the real-words conditions in Expt 1 (which did not include Jabberwocky conditions). Responses in all regions increase with constituent length. **C.** Estimated response to each condition of the real-words conditions (replicating Expt 1), the Jabberwocky conditions, and the non-constituents conditions in Expt 2. Responses in all regions increase with constituent length in the real-word conditions, and responses in all regions but LTPJ increase with constituent length in the Jabberwocky and non-constituent conditions. **D.** Estimated response to each condition of both the 24-word and 30-word items of Expt 3, both which consisted of contiguous real-word chunks that generally did not form syntactic constituents. Responses in all regions increase as a function of constituent length to a similar degree to the real-word conditions of Expts 1 and 2. **E.** Key contrasts by fROI (left-to-right): overall lexicality effect (increase in response for real-word over Jabberwocky conditions in Expt 2, averaging over length); constituent-length effect for real-word conditions in Expt 1 (slope of the line by participant from **B**); constituent-length effect for real-word conditions in

Expt 2 (slope of the red line by participant from **C**); constituent-length effect for Jabberwocky conditions in Expt 2 (slope of the blue line by participant from **C**); increase in constituent-length effect in real-word conditions over Jabberwocky in Expt 2 (difference between the slopes of the red and blue lines by participant from **C**). Starred bars indicate statistically significant effects by likelihood ratio test (corrected for false discovery rate across fROIs; (48)). Error bars show standard error of the mean over participants.

| Contrast | Expt | fROI | β | σ(β) | t | *p* |
|---|---|---|---|---|---|---|
| Constituent length for real-word conditions | 1 | **Overall** | 0.18 | 0.03 | 6.12 | < 0.001*** |
| | 1 | **LIFGorb** | 0.26 | 0.05 | 5.48 | < 0.001*** |
| | 1 | **LIFGtri** | 0.24 | 0.05 | 4.86 | 0.001** |
| | 1 | **LTP** | 0.13 | 0.03 | 4.64 | 0.001** |
| | 1 | **LaSTS** | 0.14 | 0.03 | 4.23 | 0.002** |
| | 1 | **LpSTS** | 0.20 | 0.04 | 5.52 | < 0.001*** |
| | 1 | **LTPJ** | 0.12 | 0.03 | 4.26 | 0.002** |
| Constituent length for real-word conditions | 2 | **Overall** | 0.15 | 0.02 | 6.28 | < 0.001*** |
| | 2 | **LIFGorb** | 0.21 | 0.03 | 6.44 | < 0.001*** |
| | 2 | **LIFGtri** | 0.20 | 0.04 | 5.80 | < 0.001*** |
| | 2 | **LTP** | 0.12 | 0.02 | 6.66 | < 0.001*** |
| | 2 | **LaSTS** | 0.14 | 0.02 | 7.04 | < 0.001*** |
| | 2 | **LpSTS** | 0.16 | 0.02 | 6.63 | < 0.001*** |
| | 2 | **LTPJ** | 0.09 | 0.01 | 5.82 | < 0.001*** |
| Constituent length for Jabberwocky conditions | 2 | **Overall** | 0.08 | 0.02 | 3.76 | 0.003** |
| | 2 | **LIFGorb** | 0.07 | 0.02 | 3.06 | 0.012* |
| | 2 | **LIFGtri** | 0.09 | 0.02 | 3.69 | 0.003** |
| | 2 | **LTP** | 0.06 | 0.02 | 3.96 | 0.002** |
| | 2 | **LaSTS** | 0.07 | 0.02 | 4.49 | < 0.001*** |
| | 2 | **LpSTS** | 0.14 | 0.02 | 7.90 | < 0.001*** |
| | 2 | LTPJ | 0.02 | 0.01 | 1.41 | 0.389 |
| Lexicality effect (real-word > Jabberwocky) | 2 | **Overall** | 0.67 | 0.08 | 8.39 | < 0.001*** |
| | 2 | **LIFGorb** | 0.60 | 0.09 | 6.73 | < 0.001*** |
| | 2 | **LIFGtri** | 0.60 | 0.13 | 4.53 | < 0.001*** |
| | 2 | **LTP** | 0.65 | 0.08 | 7.62 | < 0.001*** |
| | 2 | **LaSTS** | 0.76 | 0.07 | 10.95 | < 0.001*** |
| | 2 | **LpSTS** | 0.89 | 0.11 | 8.27 | < 0.001*** |
| | 2 | **LTPJ** | 0.51 | 0.07 | 7.58 | < 0.001*** |
| Constituent-length by stimulus type (real-word vs. Jabberwocky) interaction | 2 | **Overall** | 0.08 | 0.02 | 3.34 | 0.005** |
| | 2 | **LIFGorb** | 0.14 | 0.03 | 4.44 | < 0.001*** |
| | 2 | **LIFGtri** | 0.11 | 0.03 | 3.46 | 0.006** |
| | 2 | **LTP** | 0.06 | 0.02 | 2.67 | 0.033* |
| | 2 | **LaSTS** | 0.07 | 0.02 | 3.19 | 0.010* |
| | 2 | LpSTS | 0.01 | 0.03 | 0.51 | 1.000 |
| | 2 | **LTPJ** | 0.07 | 0.02 | 4.47 | < 0.001*** |
| Length effect in Expt 3 (mostly non-constituents) | 3 | **Overall** | 0.18 | 0.03 | 5.53 | < 0.001*** |
| | 3 | **LIFGorb** | 0.19 | 0.03 | 7.35 | < 0.001*** |
| | 3 | **LIFGtri** | 0.25 | 0.04 | 6.80 | < 0.001*** |
| | 3 | **LTP** | 0.17 | 0.03 | 6.52 | < 0.001*** |
| | 3 | **LaSTS** | 0.18 | 0.02 | 8.03 | < 0.001*** |
| | 3 | **LpSTS** | 0.23 | 0.03 | 7.92 | < 0.001*** |
| | 3 | **LTPJ** | 0.06 | 0.02 | 2.76 | 0.030* |

| | | | | | | |
|---|---|---|---|---|---|---|
| Length effect in Expt 1 (constituents) vs. Expt 3 (mostly non-constituents) | 1 & 3 | Overall | 0.03 | 0.03 | 0.92 | 0.733 |
| | 1 & 3 | LIFGorb | -0.02 | 0.05 | -0.44 | 1.000 |
| | 1 & 3 | LIFGtri | 0.04 | 0.06 | 0.76 | 1.000 |
| | 1 & 3 | LTP | 0.05 | 0.03 | 1.70 | 0.864 |
| | 1 & 3 | LaSTS | 0.04 | 0.03 | 1.22 | 0.864 |
| | 1 & 3 | LpSTS | 0.07 | 0.04 | 1.82 | 1.000 |
| | 1 & 3 | LTPJ | -0.03 | 0.03 | -1.00 | 0.864 |
| Length effect in Expt 2 (constituents) vs. Expt 3 (mostly non-constituents) | 2 & 3 | Overall | 0.03 | 0.03 | 0.92 | 0.362 |
| | 2 & 3 | LIFGorb | -0.02 | 0.05 | -0.44 | 1.000 |
| | 2 & 3 | LIFGtri | 0.04 | 0.06 | 0.76 | 1.000 |
| | 2 & 3 | LTP | 0.05 | 0.03 | 1.70 | 0.697 |
| | 2 & 3 | LaSTS | 0.04 | 0.03 | 1.22 | 1.000 |
| | 2 & 3 | LpSTS | 0.07 | 0.04 | 1.82 | 0.697 |
| | 2 & 3 | LTPJ | -0.03 | 0.03 | -1.00 | 1.000 |

**Table S2:** Reanalysis using PDD's anatomical ROIs as localizer masks, rather than the standard language localizer masks (parallels **Table S2**). Significance tests of key contrasts with estimates, standard errors, and *t*-values (respectively columns β, σ(β), and t). *p*-values are generated by likelihood ratio tests of linear mixed-effects models, with by-fROI results corrected for false discovery rate (FDR) applied over all six fROIs using the Benjamini-Yekutieli procedure (48) using a nominal significance level of α = 0.05. Starred *p*-values indicate statistical significance under FDR correction (*: p ≤ 0.05, **: p ≤ 0.01, *** p: ≤ 0.001). Significant regions are shown in **bold** in the fROI column.

### SI Section 10: A Comparison of the Overlapping Sets of Conditions between Our Earlier Work (Fedorenko et al., 2010) and Experiment 2 in the Current Article

PDD conditions c12, c01, jab-c12, and jab-c01 correspond respectively to the sentence (S), word list (W), Jabberwocky (J), and nonword list (N) conditions that have been investigated in several prior studies, including by our group (14). As shown in **Figure S4**, the pattern that we observed in Experiment 2 in the current study for this subset of conditions is remarkably similar to the patterns reported for Experiments 1 and 2 in ref. (14) (note that the difference in the overall response between the three experiments is most likely due to the fact that Experiment 1 in (14) and the current experiment used 12-word/nonword-long materials, and Experiment 2 in (14) used 8-word/nonword-long materials). Current Experiment 2 therefore constitutes a third within-lab replication—all with different sets of materials and non-overlapping sets of participants—of the pattern whereby sentences elicit the strongest response, word lists and Jabberwocky sentences intermediate response, and nonword lists the lowest response (see e.g., ref. (81) for another fMRI replication; see (82) for a replication in ECoG). As discussed elsewhere, including in the main text, this pattern suggests that all the regions of the language network support both the processing of word meanings and combinatorial structure building.

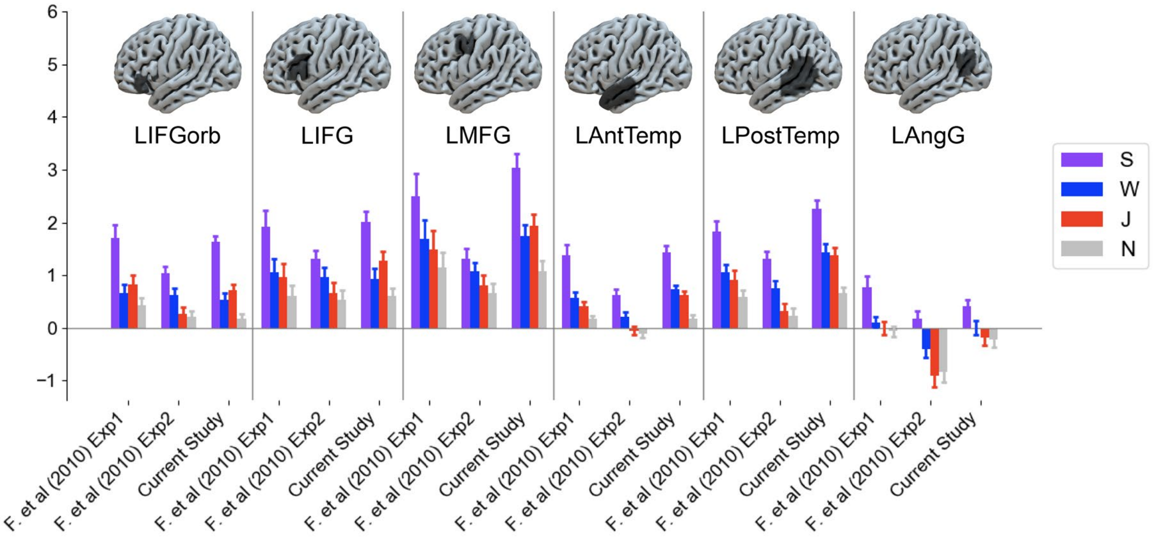

**Figure S4.** Effect estimates from the sentence (S), word list (W), Jabberwocky sentence (J), and nonword list (N) conditions from Experiments 1 and 2 of Fedorenko et al. (2010) (left and center) vs. the equivalent conditions (S=c12, W=c01, J=jab-c12, N=jab-c01) from the current Experiment 2. Error bars show standard error of the mean across participants.

### SI Section 11: Analysis of Right Hemisphere Homotopic Regions

We have thus far followed PDD in exclusively analyzing left hemisphere (LH) language regions. In light of growing interest in the contribution of the right hemisphere to language processing (83), in this section we include exploratory analyses of the key patterns within the right hemisphere (RH) homotopic language regions. Following e.g., ref. (84), we define these regions by first projecting the mirror images of our LH anatomical masks onto the right hemisphere and then following the same functional localization procedure used in the main analyses (i.e., selecting the top 10% most responsive voxels to the *sentences* > *nonwords* contrast during the localizer task). This approach allows asymmetric patterns of activation across hemispheres at the individual level while continuing to ensure functionally comparable regions of interest both within individuals (between hemispheres) and between individuals.

In direct between-hemisphere comparisons, we find significantly reduced length effects in all three experiments in the network overall and in all 6 regions in the right hemisphere relative to the left hemisphere, except in the AngG language regions in Experiment 3. We further find significantly reduced length effects for the Jabberwocky conditions of Experiment 2 in the network overall and in all individual regions except the AngG language regions. The length-driven patterns of activation are thus greatly attenuated in RH relative to LH (see **Tables S3 and S4** for full testing results). Nonetheless, as shown in **Figure S3**, the RH homotopic language areas also tend to show length effects in real-word conditions, albeit substantially weaker than those found in the left hemisphere (**Figure 3** of the main article), with fewer of these effects reaching significance (e.g., the real-word length effect is weak and not significant in RMFG, whereas it is strong and significant in LMFG). The length effect for Jabberwocky is significant only in the RH temporal language areas, indicating a generally reduced engagement of RH areas in the processing of syntactically well-formed but meaningless stimuli, relative to their LH homotopes. The RH language homotopes thus seem to show similar but attenuated patterns of response to parametric variation of the length of linguistic context.

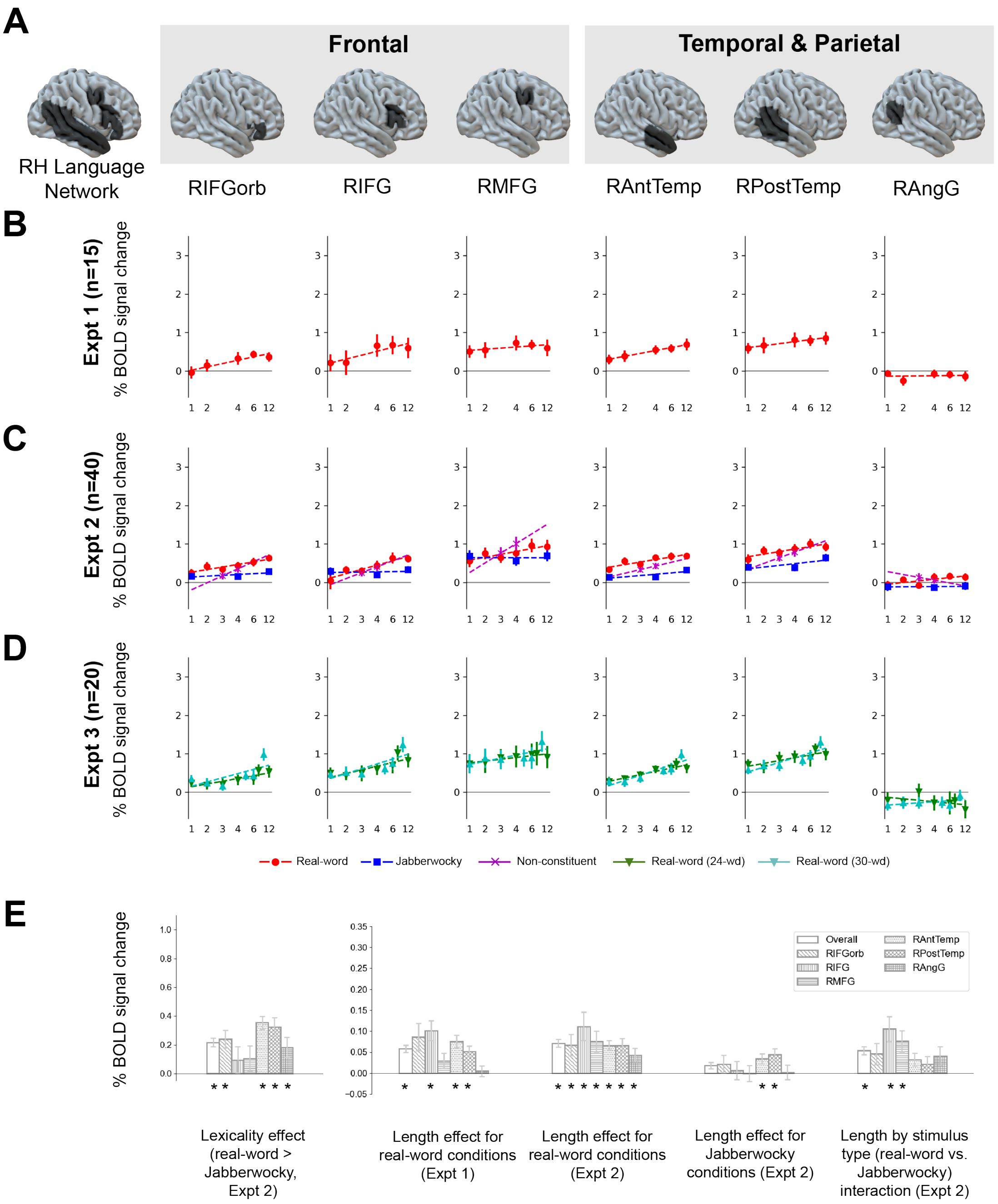

**Figure S3.** Main results (parallel of **Figure 3** of the main article) using PDD's anatomical ROIs as localizer masks, rather than the standard localizer masks. **A.** Group masks bounding the six right-hemisphere language regions. The top 10% of language-selective voxels are selected within each mask in each participant. **B.** Estimated response to each condition of the real-word conditions in Expt 1 (which did not include Jabberwocky conditions). Responses in all regions but RMFG and RAngG increase with constituent length, albeit more weakly than in their left-hemisphere homotopes (**Figure 3**). **C.** Estimated response to each condition of the real-word conditions (replicating Expt 1), the Jabberwocky conditions, and the non-constituents conditions in Expt 2. Responses in all regions increase with constituent length in the real-word conditions and with constituent length in the non-constituent conditions in all regions but RAngG, but only in temporal regions do responses increase with constituent length in the Jabberwocky conditions. **D.** Estimated response to each condition of both the 24-word and 30-word items of Expt 3, both which consisted of contiguous real-word chunks that generally did not form syntactic constituents. Responses in all regions but RAngG increase as a function of constituent length to a similar degree to the real-word conditions of Expts 1 and 2. **E.** Key contrasts by fROI (left-to-right): overall lexicality effect (increase in response for real-word over Jabberwocky conditions in Expt 2, averaging over length); constituent-length effect for real-word conditions in Expt 1 (slope of the line by participant from

**B**); constituent-length effect for real-word conditions in Expt 2 (slope of the red line by participant from **C**); constituent-length effect for Jabberwocky conditions in Expt 2 (slope of the blue line by participant from **C**); increase in constituent-length effect in real-word conditions over Jabberwocky in Expt 2 (difference between the slopes of the red and blue lines by participant from **C**). Starred bars indicate statistically significant effects by likelihood ratio test (corrected for false discovery rate across fROIs; (48)). Error bars show standard error of the mean over participants.

| **Contrast** | **Expt** | **fROI** | **β** | **σ(β)** | **t** | ***p*** |
|---|---|---|---|---|---|---|
| Constituent length for real-word conditions | 1 | **Overall** | 0.06 | 0.02 | 3.53 | 0.006** |
| | 1 | LIFGorb | 0.09 | 0.03 | 2.69 | 0.065 |
| | 1 | **LIFG** | 0.10 | 0.02 | 4.23 | 0.006** |
| | 1 | LMFG | 0.03 | 0.02 | 1.62 | 0.375 |
| | 1 | **LAntTemp** | 0.08 | 0.02 | 4.96 | 0.003** |
| | 1 | **LPostTemp** | 0.05 | 0.01 | 3.94 | 0.007** |
| | 1 | LAngG | 0.00 | 0.01 | 0.39 | 1.000 |
| Constituent length for real-word conditions | 2 | **Overall** | 0.07 | 0.02 | 3.89 | < 0.001*** |
| | 2 | **LIFGorb** | 0.07 | 0.03 | 2.66 | 0.028* |
| | 2 | **LIFG** | 0.11 | 0.03 | 3.23 | 0.012* |
| | 2 | **LMFG** | 0.08 | 0.02 | 3.01 | 0.017* |
| | 2 | **LAntTemp** | 0.07 | 0.01 | 5.74 | < 0.001*** |
| | 2 | **LPostTemp** | 0.07 | 0.02 | 3.79 | 0.004** |
| | 2 | **LAngG** | 0.04 | 0.02 | 2.66 | 0.028* |
| Constituent length for Jabberwocky conditions | 2 | Overall | 0.02 | 0.01 | 1.26 | 0.214 |
| | 2 | LIFGorb | 0.02 | 0.02 | 0.98 | 1.000 |
| | 2 | LIFG | 0.01 | 0.02 | 0.29 | 1.000 |
| | 2 | LMFG | 0.00 | 0.02 | -0.05 | 1.000 |
| | 2 | **LAntTemp** | 0.03 | 0.01 | 2.89 | 0.046* |
| | 2 | **LPostTemp** | 0.04 | 0.01 | 3.16 | 0.045* |
| | 2 | LAngG | 0.00 | 0.02 | 0.13 | 1.000 |
| Lexicality effect (real-word > Jabberwocky) | 2 | **Overall** | 0.22 | 0.07 | 3.29 | 0.004** |
| | 2 | **LIFGorb** | 0.24 | 0.06 | 3.80 | 0.002** |
| | 2 | LIFG | 0.09 | 0.09 | 1.02 | 0.773 |
| | 2 | LMFG | 0.10 | 0.09 | 1.17 | 0.737 |
| | 2 | **LAntTemp** | 0.35 | 0.04 | 7.88 | < 0.001*** |
| | 2 | **LPostTemp** | 0.32 | 0.06 | 5.00 | < 0.001*** |
| | 2 | **LAngG** | 0.18 | 0.07 | 2.59 | 0.049* |
| Constituent-length by stimulus type (real-word vs. Jabberwocky) interaction | 2 | **Overall** | 0.05 | 0.02 | 2.78 | 0.011* |
| | 2 | LIFGorb | 0.05 | 0.03 | 1.78 | 0.242 |
| | 2 | **LIFG** | 0.11 | 0.03 | 3.50 | 0.017* |
| | 2 | **LMFG** | 0.08 | 0.03 | 2.97 | 0.038* |
| | 2 | LAntTemp | 0.03 | 0.02 | 2.07 | 0.219 |
| | 2 | LPostTemp | 0.02 | 0.02 | 1.09 | 0.694 |
| | 2 | LAngG | 0.04 | 0.02 | 1.83 | 0.242 |
| Length effect in Expt 3 (mostly non-constituents) | 3 | **Overall** | 0.08 | 0.02 | 3.43 | 0.005** |
| | 3 | **LIFGorb** | 0.09 | 0.02 | 3.69 | 0.006** |
| | 3 | **LIFG** | 0.11 | 0.02 | 4.62 | < 0.001*** |
| | 3 | LMFG | 0.06 | 0.03 | 2.14 | 0.133 |
| | 3 | **LAntTemp** | 0.11 | 0.02 | 6.81 | < 0.001*** |
| | 3 | **LPostTemp** | 0.10 | 0.02 | 5.85 | < 0.001*** |
| | 3 | LAngG | -0.01 | 0.02 | -0.33 | 1.000 |

**Table S3:** Significance tests of key contrasts **in the right-hemisphere language homotopes** with estimates, standard errors, and *t*-values (respectively columns β, σ(β), and t). *p*-values are generated by likelihood ratio tests of linear mixed-effects models, with by-fROI results corrected for false discovery rate (FDR) applied over all six fROIs using the Benjamini-Yekutieli procedure (48) using a nominal significance level of $\alpha = 0.05$. Starred *p*-values indicate statistical significance under FDR correction (*: $p \leq 0.05$, **: $p \leq 0.01$, *** p: $\leq 0.001$). Significant regions are shown in **bold** in the fROI column.

| Contrast | Expt | fROI | β | σ(β) | t | *p* |
|---|---|---|---|---|---|---|
| Laterality difference of constituent length for real-word conditions | 1 | **Overall** | 0.14 | 0.03 | 4.32 | < 0.001*** |
| | 1 | **LIFGorb** | 0.20 | 0.05 | 3.71 | 0.011* |
| | 1 | **LIFG** | 0.16 | 0.04 | 3.81 | 0.011* |
| | 1 | **LMFG** | 0.16 | 0.05 | 3.42 | 0.015* |
| | 1 | **LAntTemp** | 0.07 | 0.02 | 3.19 | 0.019* |
| | 1 | **LPostTemp** | 0.15 | 0.03 | 5.75 | < 0.001*** |
| | 1 | **LAngG** | 0.08 | 0.03 | 3.08 | 0.020* |
| Laterality difference of constituent length for real-word conditions | 2 | **Overall** | 0.11 | 0.02 | 4.96 | < 0.001*** |
| | 2 | **LIFGorb** | 0.17 | 0.03 | 6.50 | < 0.001*** |
| | 2 | **LIFG** | 0.12 | 0.03 | 3.74 | 0.002** |
| | 2 | **LMFG** | 0.17 | 0.02 | 7.00 | < 0.001*** |
| | 2 | **LAntTemp** | 0.07 | 0.02 | 4.67 | < 0.001*** |
| | 2 | **LPostTemp** | 0.09 | 0.01 | 6.30 | < 0.001*** |
| | 2 | **LAngG** | 0.05 | 0.02 | 2.81 | 0.019* |
| Laterality difference of constituent length for Jabberwocky conditions | 2 | **Overall** | 0.09 | 0.03 | 3.50 | 0.007** |
| | 2 | **LIFGorb** | 0.08 | 0.02 | 3.99 | < 0.001*** |
| | 2 | **LIFG** | 0.13 | 0.02 | 5.50 | < 0.001*** |
| | 2 | **LMFG** | 0.18 | 0.03 | 6.61 | < 0.001*** |
| | 2 | **LAntTemp** | 0.05 | 0.01 | 3.98 | < 0.001*** |
| | 2 | **LPostTemp** | 0.09 | 0.01 | 6.38 | < 0.001*** |
| | 2 | LAngG | 0.00 | 0.02 | 0.05 | 1.000 |
| Laterality difference of lexicality effect (real-word > Jabberwocky) | 2 | **Overall** | 0.53 | 0.08 | 6.46 | < 0.001*** |
| | 2 | **LIFGorb** | 0.44 | 0.09 | 5.05 | < 0.001*** |
| | 2 | **LIFG** | 0.59 | 0.11 | 5.22 | < 0.001*** |
| | 2 | **LMFG** | 0.80 | 0.09 | 9.01 | < 0.001*** |
| | 2 | **LAntTemp** | 0.42 | 0.06 | 6.67 | < 0.001*** |
| | 2 | **LPostTemp** | 0.62 | 0.08 | 8.20 | < 0.001*** |
| | 2 | **LAngG** | 0.31 | 0.07 | 4.19 | < 0.001*** |
| Laterality difference of constituent-length by stimulus type (real-word vs. Jabberwocky) interaction | 2 | Overall | 0.02 | 0.02 | 1.42 | 0.173 |
| | 2 | LIFGorb | 0.08 | 0.03 | 2.97 | 0.075 |
| | 2 | LIFG | 0.00 | 0.03 | -0.10 | 1.000 |
| | 2 | LMFG | -0.01 | 0.03 | -0.23 | 1.000 |
| | 2 | LAntTemp | 0.02 | 0.02 | 1.20 | 1.000 |
| | 2 | LPostTemp | 0.00 | 0.02 | -0.05 | 1.000 |
| | 2 | LAngG | 0.05 | 0.03 | 1.81 | 0.569 |
| Laterality difference of length effect in Expt 3 (mostly non-constituents) | 3 | **Overall** | 0.12 | 0.02 | 5.59 | < 0.001*** |
| | 3 | **LIFGorb** | 0.13 | 0.03 | 5.07 | < 0.001*** |
| | 3 | **LIFG** | 0.15 | 0.03 | 5.04 | < 0.001*** |
| | 3 | **LMFG** | 0.12 | 0.03 | 4.46 | < 0.001*** |
| | 3 | **LAntTemp** | 0.10 | 0.02 | 5.35 | < 0.001*** |
| | 3 | **LPostTemp** | 0.13 | 0.02 | 5.92 | < 0.001*** |
| | 3 | LAngG | 0.07 | 0.03 | 2.52 | 0.051 |

**Table S4:** Significance tests of **laterality difference (LH – RH)** of key contrasts with estimates, standard errors, and *t*-values (respectively columns β, σ(β), and t). *p*-values are generated by likelihood ratio tests of linear mixed-effects models, with by-fROI results corrected for false discovery rate (FDR) applied over all six fROIs using the Benjamini-Yekutieli procedure (48) using a nominal significance level of $\alpha = 0.05$. Starred *p*-values indicate statistical significance under FDR correction (*: p ≤ 0.05, **: p ≤ 0.01, *** p: ≤ 0.001). Significant regions are shown in **bold** in the fROI column.

### References

1. C. Pallier, A.-D. Devauchelle, S. Dehaene, Cortical representation of the constituent structure of sentences. *Proc. Natl. Acad. Sci.* **108**, 2522–2527 (2011).
2. J. A. Fodor, *Modularity of Mind* (MIT Press, 1983).
3. S. Dehaene, F. Meyniel, C. Wacongne, L. Wang, C. Pallier, The neural representation of sequences: from transition probabilities to algebraic patterns and linguistic trees. *Neuron* **88**, 2–19 (2015).
4. S. Dehaene, The Demodularization Hypothesis. *The Neocortex* **27**, 293 (2019).
5. G. Kempen, Prolegomena to a neurocomputational architecture for human grammatical encoding and decoding. *Neuroinformatics* **12**, 111–142 (2014).
6. I. Hertrich, S. Dietrich, H. Ackermann, The role of the supplementary motor area for speech and language processing. *Neurosci. Biobehav. Rev.* **68**, 602–610 (2016).
7. M. J. Nelson, *et al.*, Neurophysiological dynamics of phrase-structure building during sentence processing. *Proc. Natl. Acad. Sci.* **114**, E3669--E3678 (2017).
8. L. Wang, L. Uhrig, B. Jarraya, S. Dehaene, Representation of numerical and sequential patterns in macaque and human brains. *Curr. Biol.* **25**, 1966–1974 (2015).
9. C. Pattamadilok, S. Dehaene, C. Pallier, A role for left inferior frontal and posterior superior temporal cortex in extracting a syntactic tree from a sentence. *cortex* **75**, 44–55 (2016).
10. S. Dehaene, F. Al Roumi, Y. Lakretz, S. Planton, M. Sablé-Meyer, Symbols and mental programs: a hypothesis about human singularity. *Trends Cogn. Sci.* (2022).
11. S. R. Hage, A. Nieder, Dual neural network model for the evolution of speech and language. *Trends Neurosci.* **39**, 813–829 (2016).
12. B. M. Mazoyer, *et al.*, The cortical representation of speech. *J. Cogn. Neurosci.* **5**, 467–479 (1993).
13. C. Humphries, J. R. Binder, D. A. Medler, E. Liebenthal, Syntactic and semantic modulation of neural activity during auditory sentence comprehension. *J. Cogn. Neurosci.* **18**, 665–679 (2006).
14. E. Fedorenko, P.-J. Hsieh, A. Nieto-Castañón, S. Whitfield-Gabrieli, N. Kanwisher, New method for fMRI investigations of language: defining ROIs functionally in individual subjects. *J. Neurophysiol.* **104**, 1177–1194 (2010).
15. K. Mahowald, E. Fedorenko, Reliable individual-level neural markers of high-level language processing: A necessary precursor for relating neural variability to behavioral and genetic variability. *Neuroimage* **139**, 74–93 (2016).
16. W. Matchin, C. Hammerly, E. Lau, The role of the IFG and pSTS in syntactic prediction: Evidence from a parametric study of hierarchical structure in fMRI. *Cortex* **88**, 106–123 (2017).
17. R. C. Berwick, G. J. L. Beckers, K. Okanoya, J. J. Bolhuis, A bird's eye view of human language evolution. *Front. Evol. Neurosci.* **4**, 5 (2012).
18. S. F. Cappa, Imaging semantics and syntax. *Neuroimage* **61**, 427–431 (2012).
19. W. T. Fitch, Toward a computational framework for cognitive biology: unifying approaches from cognitive neuroscience and comparative cognition. *Phys. Life Rev.* **11**, 329–364 (2014).
20. W. T. Fitch, M. D. Martins, Hierarchical processing in music, language, and action: Lashley revisited. *Ann. N. Y. Acad. Sci.* **1316**, 87–104 (2014).
21. K. Friston, G. Buzsáki, The functional anatomy of time: what and when in the brain. *Trends Cogn. Sci.* **20**, 500–511 (2016).

22. I. Bornkessel-Schlesewsky, M. Schlesewsky, Reconciling time, space and function: a new dorsal--ventral stream model of sentence comprehension. *Brain Lang.* **125**, 60–76 (2013).

23. I. Bornkessel-Schlesewsky, M. Schlesewsky, S. L. Small, J. P. Rauschecker, Neurobiological roots of language in primate audition: Common computational properties. *Trends Cogn. Sci.* **19**, 142–150 (2015).

24. M. A. Skeide, J. Brauer, A. D. Friederici, Brain functional and structural predictors of language performance. *Cereb. Cortex* **26**, 2127–2139 (2016).

25. E. Zaccarella, A. D. Friederici, Merge in the human brain: A sub-region based functional investigation in the left pars opercularis. *Front. Psychol.* **6**, 1818 (2015).

26. E. Zaccarella, M. Schell, A. D. Friederici, Reviewing the functional basis of the syntactic Merge mechanism for language: A coordinate-based activation likelihood estimation meta-analysis. *Neurosci. Biobehav. Rev.* **80**, 646–656 (2017).

27. S. M. Wilson, *et al.*, What role does the anterior temporal lobe play in sentence-level processing? Neural correlates of syntactic processing in semantic variant primary progressive aphasia. *J. Cogn. Neurosci.* **26**, 970–985 (2014).

28. S. M. Frankland, J. D. Greene, Concepts and compositionality: in search of the brain's language of thought. *Annu. Rev. Psychol.* **71**, 273–303 (2020).

29. A. Bautista, S. M. Wilson, Neural responses to grammatically and lexically degraded speech. *Lang. Cogn. Neurosci.* **31**, 567–574 (2016).

30. K. J. Friston, *et al.*, Spatial registration and normalization of images. *Hum. Brain Mapp.* **3**, 165–189 (1995).

31. A. Nieto-Castanon, *Handbook of functional connectivity Magnetic Resonance Imaging methods in CONN* (Hilbert Press, 2020).

32. J. Ashburner, K. J. Friston, Unified segmentation. *Neuroimage* **26**, 839–851 (2005).

33. R. M. Braga, L. M. DiNicola, H. C. Becker, R. L. Buckner, Situating the left-lateralized language network in the broader organization of multiple specialized large-scale distributed networks. *J. Neurophysiol.* **124**, 1415–1448 (2020).

34. B. Lipkin, *et al.*, Probabilistic atlas for the language network based on precision fMRI data from >800 individuals. *Sci. data* **9** (2022).

35. E. Fedorenko, S. L. Thompson-Schill, Reworking the language network. *Trends Cogn. Sci.* **18**, 120–126 (2014).

36. T. L. Scott, J. Gallée, E. Fedorenko, A new fun and robust version of an fMRI localizer for the frontotemporal language system. *Cogn. Neurosci.* **8**, 167–176 (2017).

37. X. Chen, *et al.*, The human language system does not support music processing. *bioRxiv* (2021).

38. S. Malik-Moraleda, *et al.*, The universal language network: A cross-linguistic investigation spanning 45 languages and 11 language families. *bioRxiv* (2022).

39. A. A. Ivanova, *et al.*, The Effect of Task on Brain Activity during Sentence Processing in *12th Annual Meeting of the Society for the Neurobiology of Language (SNL20)*, (2020).

40. I. Blank, Z. Balewski, K. Mahowald, E. Fedorenko, Syntactic processing is distributed across the language system. *Neuroimage* **127**, 307–323 (2016).

41. I. Blank, N. Kanwisher, E. Fedorenko, A functional dissociation between language and multiple-demand systems revealed in patterns of BOLD signal fluctuations. *J. Neurophysiol.* **112**, 1105–1118 (2014).

42. Y. Tie, *et al.*, Defining language networks from resting-state fMRI for surgical planning -- A feasibility study. *Hum. Brain Mapp.* **35**, 1018–1030 (2014).

43. P. Branco, D. Seixas, S. L. Castro, Mapping language with resting-state functional magnetic resonance imaging: A study on the functional profile of the language network. *Hum. Brain Mapp.* **41**, 545–560 (2020).

44. M. F. Glasser, *et al.*, A multi-modal parcellation of human cerebral cortex. *Nature* **536**, 171–178 (2016).

45. A. Nieto-Castañón, E. Fedorenko, Subject-specific functional localizers increase sensitivity and functional resolution of multi-subject analyses. *Neuroimage* **63**, 1646–1669 (2012).

46. N. Kriegeskorte, W. K. Simmons, P. S. F. Bellgowan, C. I. Baker, Circular analysis in systems neuroscience: The dangers of double dipping. *Nat. Neurosci.* **12**, 535–540 (2009).

47. E. Fedorenko, M. K. Behr, N. Kanwisher, Functional specificity for high-level linguistic processing in the human brain. *Proc. Natl. Acad. Sci.* **108**, 16428–16433 (2011).

48. Y. Benjamini, D. Yekutieli, The control of the false discovery rate in multiple testing under dependency. *Ann. Stat.* **29**, 1165–1188 (2001).

49. M. P. Marcus, B. Santorini, M. A. Marcinkiewicz, Building a large annotated corpus of English: the Penn Treebank. *Comput. Linguist.* **19**, 313–330 (1993).

50. R. Futrell, *et al.*, The Natural Stories corpus: a reading-time corpus of English texts containing rare syntactic constructions. *Lang. Resour. Eval.*, 1–15 (2020).

51. E. Keuleers, M. Brysbaert, Wuggy: A multilingual pseudoword generator. *Behav. Res. Methods* **42**, 627–633 (2010).

52. J. Nivre, *et al.*, Universal Dependencies v1: A Multilingual Treebank Collection in *LREC*, (2016).

53. N. Chomsky, *The minimalist program* (MIT Press, 1995).

54. L. Nguyen, M. van Schijndel, W. Schuler, Accurate Unbounded Dependency Recovery using Generalized Categorial Grammars in *Proceedings of COLING 2012*, (2012).

55. M. van Schijndel, A. Exley, W. Schuler, A model of language processing as hierarchic sequential prediction. *Top. Cogn. Sci.* **5**, 522–540 (2013).

56. J. Hale, Uncertainty about the rest of the sentence. *Cogn. Sci.* **30**, 609–642 (2006).

57. J. Brennan, *et al.*, Syntactic structure building in the anterior temporal lobe during natural story listening. *Brain Lang.* **120**, 163–173 (2012).

58. J. Brennan, E. P. Stabler, S. E. Van Wagenen, W.-M. Luh, J. T. Hale, Abstract linguistic structure correlates with temporal activity during naturalistic comprehension. *Brain Lang.* **157**, 81–94 (2016).

59. E. Gibson, “The Dependency Locality Theory: A distance-based theory of linguistic complexity” in *Image, Language, Brain*, A. Marantz, Y. Miyashita, W. O’Neil, Eds. (MIT Press, 2000), pp. 95–106.

60. P. C. Gordon, R. Hendrick, M. Johnson, Memory Interference during Language Processing. *J. Exp. Psychol. Learn. Mem. Cogn.* **27**, 1411–1423 (2001).

61. R. L. Lewis, S. Vasishth, An activation-based model of sentence processing as skilled memory retrieval. *Cogn. Sci.* **29**, 375–419 (2005).

62. N. E. Rasmussen, W. Schuler, Left-Corner Parsing With Distributed Associative Memory Produces Surprisal and Locality Effects. *Cogn. Sci.* **42**, 1009–1042 (2018).

63. C. Shain, I. A. Blank, E. Fedorenko, E. Gibson, W. Schuler, Robust effects of working memory

demand during naturalistic language comprehension in language-selective cortex. *J. Neurosci.* (2022) https:/doi.org/10.1523/JNEUROSCI.1894-21.2022.

64. D. J. Grodner, E. Gibson, Consequences of the serial nature of linguistic input. *Cogn. Sci.* **29**, 261–291 (2005).

65. E. Chen, E. Gibson, F. Wolf, Online syntactic storage costs in sentence comprehension. *J. Mem. Lang.* **52**, 144–169 (2005).

66. C. Shain, M. van Schijndel, R. Futrell, E. Gibson, W. Schuler, Memory access during incremental sentence processing causes reading time latency in *Proceedings of the Workshop on Computational Linguistics for Linguistic Complexity (CL4LC)*, (2016), pp. 49–58.

67. S. L. Frank, R. Bod, Insensitivity of the human sentence-processing system to hierarchical structure. *Psychol. Sci.* (2011).

68. N. J. Smith, R. Levy, The effect of word predictability on reading time is logarithmic. *Cognition* **128**, 302–319 (2013).

69. R. M. Willems, S. L. Frank, A. D. Nijhof, P. Hagoort, A. den Bosch, Prediction during natural language comprehension. *Cereb. Cortex* **26**, 2506–2516 (2015).

70. A. Goodkind, K. Bicknell, Predictive power of word surprisal for reading times is a linear function of language model quality in *Proceedings of the 8th Workshop on Cognitive Modeling and Computational Linguistics (CMCL 2018)*, (2018), pp. 10–18.

71. C. Shain, I. Blank, M. van Schijndel, W. Schuler, E. Fedorenko, fMRI reveals language-specific predictive coding during naturalistic sentence comprehension. *Neuropsychologia* **138**, 107307 (2020).

72. J. Hale, A Probabilistic Earley Parser as a Psycholinguistic Model in *Proceedings of the Second Meeting of the North American Chapter of the Association for Computational Linguistics*, (2001), pp. 159–166.

73. R. Levy, Expectation-based syntactic comprehension. *Cognition* **106**, 1126–1177 (2008).

74. K. Heafield, I. Pouzyrevsky, J. H. Clark, P. Koehn, Scalable modified Kneser-Ney language model estimation in *Proceedings of the 51st Annual Meeting of the Association for Computational Linguistics*, (2013), pp. 690–696.

75. D. Graff, C. Cieri, English Gigaword LDC2003T05 (2003).

76. V. Demberg, F. Keller, Data from eye-tracking corpora as evidence for theories of syntactic processing complexity. *Cognition* **109**, 193–210 (2008).

77. A. Lopopolo, S. L. Frank, A. den Bosch, R. M. Willems, Using stochastic language models (SLM) to map lexical, syntactic, and phonological information processing in the brain. *PLoS One* **12**, e0177794 (2017).

78. V. Fossum, R. Levy, Sequential vs. Hierarchical Syntactic Models of Human Incremental Sentence Processing in *Proceedings of {{CMCL}} 2012*, (Association for Computational Linguistics, 2012).

79. M. van Schijndel, W. Schuler, Hierarchic syntax improves reading time prediction in *Proceedings of NAACL-HLT 2015*, (Association for Computational Linguistics, 2015).

80. D. Bates, M. Mächler, B. Bolker, S. Walker, Fitting linear mixed-effects models using lme4. *J. Stat. Softw.* **67**, 1–48 (2015).

81. M. Bedny, A. Pascual-Leone, D. Dodell-Feder, E. Fedorenko, R. Saxe, Language processing in the occipital cortex of congenitally blind adults. *Proc. Natl. Acad. Sci.* **108**, 4429–4434 (2011).

82. E. Fedorenko, *et al.*, Neural correlate of the construction of sentence meaning. *Proc. Natl. Acad.

*Sci.* **113**, E6256--E6262 (2016).

83. K. C. Martin, *et al.*, A weak shadow of early life language processing persists in the right hemisphere of the mature brain. *Neurobiol. Lang.* **3**, 364–385 (2022).

84. C. Shain, A. Paunov, X. Chen, B. Lipkin, E. Fedorenko, No evidence of theory of mind reasoning in the human language network. *bioRxiv* (2022).
